# The Toxoplasma Vacuolar H^+^-ATPase regulates intracellular pH, and impacts the maturation of essential secretory proteins

**DOI:** 10.1101/515296

**Authors:** Andrew J. Stasic, Nathan M. Chasen, Eric J. Dykes, Stephen Vella, Beejan Asady, Vincent J. Starai, Silvia N.J. Moreno

**Author notes:** Corresponding author: Silvia NJ Moreno, Department of Cellular Biology, University of Georgia, 350A Coverdell Center, 500 D.W. Brooks Dr. Athens, GA 30602-7399, U.S.A.

## Abstract

Vacuolar-proton ATPases (V-H^+^-ATPases) are conserved complexes that couple the hydrolysis of ATP to the pumping of protons across membranes. V-H^+^-ATPases are known to play diverse roles in cellular physiology. We studied the *Toxoplasma gondii* V-H^+^-ATPase complex and discovered a novel dual role of the pump in protecting parasites against ionic stress and in the maturation of secretory proteins in endosomal-like compartments. *Toxoplasma* V-H^+^-ATPase subunits localize to the plasma membrane and to acidic vesicles and characterization of conditional mutants of the a1 subunit highlighted the functionality of the complex at both locations. Microneme and rhoptry proteins are required for invasion and modulation of host cells and they traffic via endosome-like compartments in which proteolytic maturation occurs. We show that the V-H^+^-ATPase supports the maturation of rhoptry and microneme proteins, and their maturases, during their traffic to their corresponding organelles. This work underscores a novel role for V-H^+^-ATPases in regulating virulence pathways.

**HIGHLIGHTS:**

- The V-H^+^-ATPase localizes to the plasma membrane and pumps protons outside the cell
- The V-H^+^-ATPase localizes to pro-rhoptries in dividing parasites and to the plant-like vacuole in extracellular tachyzoites.
- The Vacuolar-H^+^-ATPase supports the maturation of rhoptry and microneme proteins.
- Loss of V-H^+^-ATPase activity leads to hyperaccumulation of immature rhoptry and microneme proteins and defective microneme distribution and secretion.

## INTRODUCTION

*Toxoplasma gondii* is an Apicomplexan parasite that infects a wide range of hosts, including humans. *T. gondii* infections are usually asymptomatic in healthy adults, but in immunosuppressed individuals, infections can cause serious complications and be fatal. The pathogenicity of this obligate intracellular parasite is dependent on its lytic cycle, which is characterized by invasion of mammalian cells, replication inside a parasitophorous vacuole (PV), and egress (Black, M. W. and Boothroyd, J. C., 2000). As *T. gondii* progresses through its lytic cycle, the ionic composition of the surrounding environment to which it is exposed changes dramatically from low calcium, sodium and chloride and high potassium inside host cells to high calcium, sodium and chloride and low potassium in the extracellular milieu. Sophisticated regulatory mechanisms are in place for the parasite to deal with these changes and also to use these ionic gradients for its own benefit such as filling its intracellular calcium stores (Pace et al., 2014). *T. gondii* tachyzoites, the fast growing forms, replicate inside their host cell, which lyses upon exit of the parasite, a phase responsible for the pathology of *Toxoplasma*. Our laboratory has previously characterized an organelle termed plant-like vacuole (PLV or VAC) that becomes prominent in extracellular tachyzoites and fragments shortly after invasion in intracellular parasites. The PLV is thought to protect parasites against ionic and osmotic stress (Miranda et al., 2010; Parussini et al., 2010). The PLV houses lytic enzymes like cathepsins, plant-like pumps and channels like the vacuolar proton pyrophosphatase, and aquaporins that act to regulate ions and/or as a post Golgi sorting compartment for secretory proteins destined to the micronemes, rhoptries, and acidocalcisomes. The PLV plays a central role during the extracellular phase of the parasite not only in resisting environmental stress but also in preparing parasites for subsequent host cell invasion. Most of the functions of this organelle depend on its ability to maintain an acidic gradient at the membrane, which is driven by 2 distinct electrogenic proton pumps: the plant-like H^+^-translocating inorganic pyrophosphatase (H^+^-PPase) and the vacuolar-H^+^-ATPase (V-H^+^-ATPase).

The V-H^+^-ATPase is an evolutionarily conserved proton pump that couples the hydrolysis of ATP with the translocation of protons across membranes, often into the lumen of a vesicle. Typically, these protein complexes consist of at least 14 different subunits that compose a membrane anchoring **V_0_ domain** (*a, c, c’, c”, d*, and *e* subunits) and a peripheral **V_1_ domain** (subunits A to H) (Forgac, 1989). In this work, we characterize the a1 subunit of the V-H^+^-ATPase complex in detail. This subunit *a* is a 100 kDa transmembrane protein containing an amino-terminal cytoplasmic domain and a carboxy-terminal hydrophobic domain containing 8–9 transmembrane helices (Leng et al., 1999). We investigated the link between the V-H^+^ATPase and PLV function to gain knowledge of the mechanism by which this organelle protects parasites against ionic stress and in addition its role in sorting and maturation of essential secretory proteins like microneme and rhoptry proteins. Our data indicate that the V-H^+^ ATPase is also functional at the plasma membrane of *T. gondii* tachyzoites where it pumps protons out of the parasite. We propose a model for the dual role of this multipurpose pump and its adaptation to the unique needs of intracellular and extracellular tachyzoites and their parasitism.

## RESULTS

### Genomic organization and expression of the *T. gondii vha1* gene

V-H^+^-ATPases are multisubunit complexes composed of two domains, V_1_ and V_0_ (**Fig. S1A**), which couple the hydrolysis of ATP with the transport of protons across membranes (Forgac, 1999). In yeasts, the peripheral V_1_ domain consists of eight subunits (A–H) and carries out ATP hydrolysis, while the integral V_0_ domain, which comprises subunits *a, c, c’, c’’, d*, and *e*, is responsible for transporting protons to the lumen of the vacuole. Subunit *a* of the V_0_ domain is a 100-kDa integral membrane protein that spans both domains of the protein complex and is involved in complex assembly (Forgac, 1999) (**Fig. S1A**). The N-terminal domain is cytoplasmic, connects V_1_ and V_0_, and stabilizes the complex during rotary catalysis (Forgac, 1999; Wang et al., 2008). The C-terminal domain is membrane-embedded, contains eight transmembrane helices, and is involved in proton transport into the proteolipid ring (Wang et al., 2008). Homology searches in ToxoDB predicted the presence of most subunits of the V-H^+^-ATPase with the exception of *e* and *c*’’ (**Table 1**). *T. gondii* appears to express two isoforms, *a*1 and *a*2 (TgGT1_232830 and TgGT1_290720) of the *a* subunit. We chose to characterize the a1 subunit because of its central role in translocating protons and potential interaction with both domains V_0_ and V_1_. Subunit *a*1 (TgGT1_232830) had a higher BLAST expectant score as compared with the yeast Vph1p (**Table 1**), stronger evidence for expression, and a stronger phenotype score than *a*2 according to the information in ToxoDB (Sidik et al., 2016). We therefore decided to focus our current studies on the *a*1 isoform. We termed this gene *Tgvha1* (*vha1* henceforth), which encodes for a predicted protein of 909 amino acids (VHa1) with an N-terminal signal peptide encompassing the first 26 amino acids. The predicted topology of VHa1 supports the presence of 7 transmembrane domains (**Fig. S1B**).

**Table 1:**
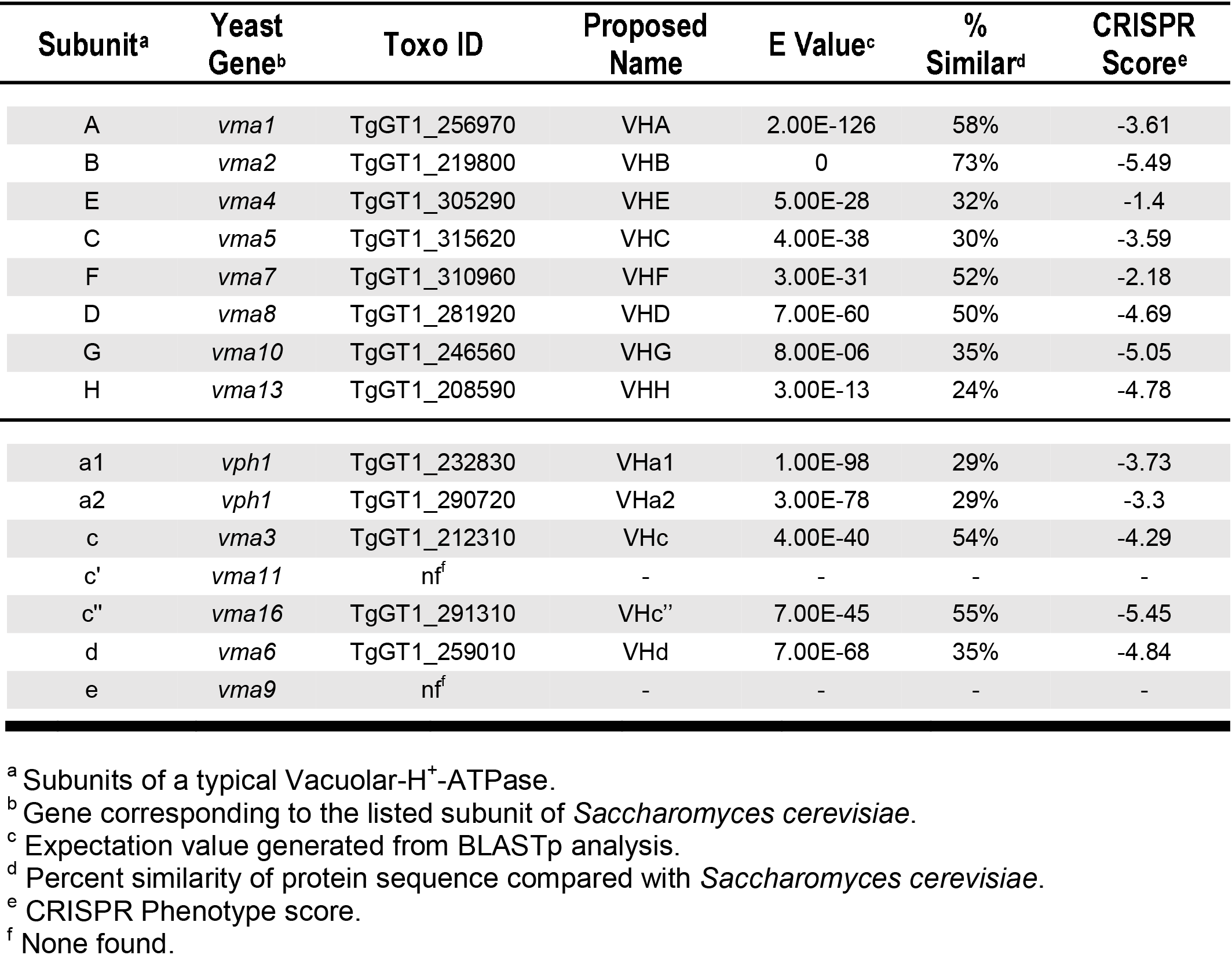
Identification of *T. gondii* V-H^+^-ATPase subunits.

|  | Subunit<sup>a</sup> | Yeast Gene<sup>b</sup> | Toxo ID | Proposed Name | E Value<sup>c</sup> | % Similar<sup>d</sup> | CRISPR Score<sup>e</sup> |
| --- | --- | --- | --- | --- | --- | --- | --- |
| V<sub>1</sub> | A | *vma1* | TgGT1_256970 | VHA | 2.00E-126 | 58% | -3.61 |
|  | B | *vma2* | TgGT1_219800 | VHB | 0 | 73% | -5.49 |
|  | E | *vma4* | TgGT1_305290 | VHE | 5.00E-28 | 32% | -1.4 |
|  | C | *vma5* | TgGT1_315620 | VHC | 4.00E-38 | 30% | -3.59 |
|  | F | *vma7* | TgGT1_310960 | VHF | 3.00E-31 | 52% | -2.18 |
|  | D | *vma8* | TgGT1_281920 | VHD | 7.00E-60 | 50% | -4.69 |
|  | G | *vma10* | TgGT1_246560 | VHG | 8.00E-06 | 35% | -5.05 |
|  | H | *vma13* | TgGT1_208590 | VHH | 3.00E-13 | 24% | -4.78 |
| V<sub>0</sub> | a1 | *vph1* | TgGT1_232830 | VHa1 | 1.00E-98 | 29% | -3.73 |
|  | a2 | *vph1* | TgGT1_290720 | VHa2 | 3.00E-78 | 29% | -3.3 |
|  | c | *vma3* | TgGT1_212310 | VHc | 4.00E-40 | 54% | -4.29 |
|  | c' | *vma11* | nf<sup>f</sup> | - | - | - | - |
|  | c'' | *vma16* | TgGT1_291310 | VHc'' | 7.00E-45 | 55% | -5.45 |
|  | d | *vma6* | TgGT1_259010 | VHd | 7.00E-68 | 35% | -4.84 |
|  | e | *vma9* | nf<sup>f</sup> | - | - | - | - |

<sup>a</sup> Subunits of a typical Vacuolar-H<sup>+</sup>-ATPase.
<sup>b</sup> Gene corresponding to the listed subunit of *Saccharomyces cerevisiae*.
<sup>c</sup> Expectation value generated from BLASTp analysis.
<sup>d</sup> Percent similarity of protein sequence compared with *Saccharomyces cerevisiae*.
<sup>e</sup> CRISPR Phenotype score.
<sup>f</sup> None found.

### The V-H^+^-ATPase localizes to the plasma membrane and the PLV

To determine the localization of the V-H^+^-ATPase in *T. gondii*, the *vha1 gene* (TgGT1_232830) was endogenously tagged with a 3xHA at the C-terminus (**Fig. 1A**) using the pLIC plasmid approach (Huynh and Carruthers, 2009; Sheiner et al., 2011); proper incorporation of this tag was confirmed by PCR (**Figs. S1C, D**). Expression of the fusion protein VHa1-3xHA was verified by western blot analysis (**Fig. 1B**) and was found to primarily localize to the plasma membrane in intracellular parasites (**Fig. 1Ci**). Extracellular parasites also showed localization to the plasma membrane as shown by IFAs with anti-HA, which labels the surface in a similar pattern to the surface marker α-SAG1 (**Fig. 1Cii**). The specific localization of VHa1-3XHA to the plasma membrane was shown by treating parasites with *Clostridium septicum* alpha-toxin, which induces separation of the plasma membrane away from the IMC (Wichroski et al., 2002). Under these conditions the IFAs with anti-HA and an anti-IMC showed that VHa1-3xHA did not colocalize with the IMC, and followed the plasma membrane (**Fig. S2A**). We also showed that VHa1-3XHA co-localized with the plasma membrane marker SAG1 (**Fig. S2B**) in intracellular parasites. We observed some intracellular labeling of undefined vesicles in intracellular parasites (Fig. 1C and S2B) but the plasma membrane labeling was the most predominant (Fig 1C). In extracellular parasites, VHa1 also localized to the plasma membrane but we also observed a large intracellular compartment that became especially prominent with time outside the host cell (**Figs. 1Cii-iv and S2C**). We performed a series of IFA co-localization studies and found that the vacuolar proton pyrophosphatase (TgVP1) and cathepsin L (TgCPL) (Parussini et al., 2010), both known markers of the plant-like vacuole (PLV) co-localized with VHa1 (Miranda et al., 2010) (**Figs. 1Ciii and iv and S2C**). Immuno-EM confirmed the presence of VHa1 at the plasma membrane and the PLV (**Fig. 1D**). The VHa1-3xHA labeled a large central structure (**see Fig. S2C**) that co-localized with the PLV marker TgVP1 shortly after egress (**Fig. S2G**). The V-H^+^-ATPase localization to the PLV became more evident and noticeable as the parasite stay extracellular and the PLV itself become more prominent. We further investigated the PLV localization by super resolution microscopy (**Figs. 1Ciii, and S2C**). In addition, we confirmed this localization by immunofluorescence analysis (IFAs) with specific antibodies generated in mice against the *a*1 subunit (**Fig. S2I**). Western blots of *Toxoplasma* lysates developed with the affinity purified mouse serum showed a band corresponding to the correct size of VHa1 and IFA’s using anti-VHa1 showed similar localization in extracellular parasites as VHa1-HA (**Figs. S2H, I**). These results confirmed that the *T. gondii* V-H^+^-ATPase localizes to the plasma membrane and the PLV. The traffic of the V-H^+^-ATPase to the PLV and its underlying mechanism remains to be characterized. We do not believe that it is being turned over or degraded because we did not observe degradation products in immunoblots (**Fig. S2H**) and our data indicates that the complex is functional at the PLV.

**FIGURE 1:**
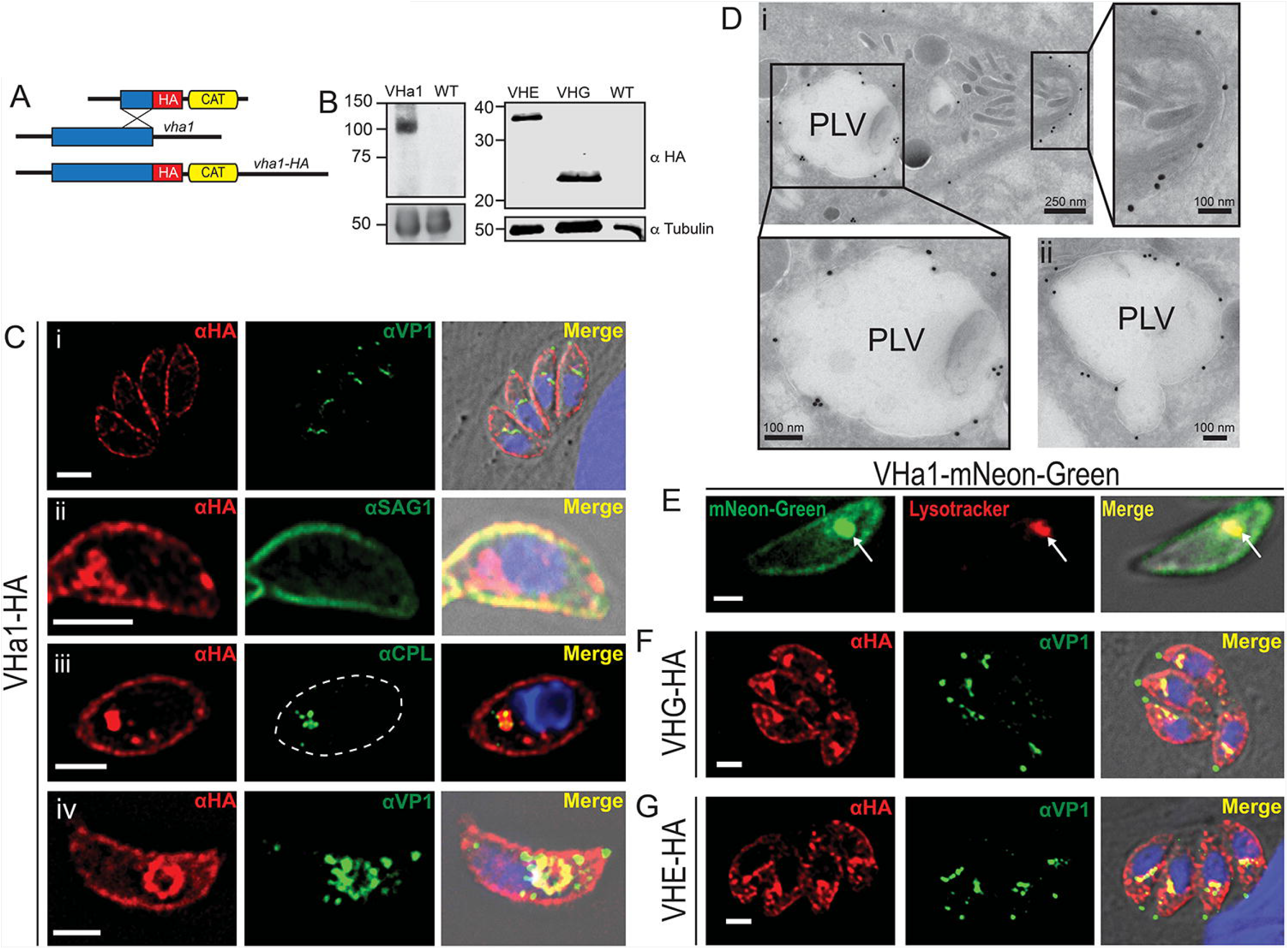
VHa1 localizes to the plasma membrane and the PLV. A) Cartoon showing the 3xHA tagging strategy. HA, hemagglutinin; CAT, chloramphenicol acetyltransferase. B) Western blots analysis of lysates of parasites expressing *vha1* C-terminally tagged with 3xHA (left panel). Lysates of tachyzoites expressing subunits E and G tagged with 3xHA are shown in the right panel. The sizes of the bands agree with the predicted MW of the three subunits: 106 kDa, 39 kDa, and 22 kDa respectively. C) Immunofluorescence assays (IFA) with VHa1-HA showing i) the localization of VHa1 in intracellular tachyzoites; ii) co-localization with SAG1 antibody; iii) super resolution IFA’s showing co-localization with the PLV marker CPL. Dashed line marks the shape of the parasite; iv) extracellular tachyzoites showing co-localization with the PLV marker VP1. D) Representative immuno-EM with anti-HA antibodies showing labeling at the i) plasma membrane and PLV. ii) Labeling of the PLV just after fusing with a vesicle. E) VHa1-mNeon-Green tagged clones showing co-localization with lysotracker red. F) IFA of VHG-HA in intracellular tachyzoites with PLV marker VP1. G) IFA of VHE-HA in intracellular tachyzoites with PLV marker VP1. The thresholded Mander’s colocalization coefficient (on a scale from −1 to 1 where 1 is perfect colocalization) of VP1/VHa1 and CPL/VHa1 of extracellular tachyzoites from 3 independent trials were 0.33 and 0.62, respectively. All IFA scale bars are 2 μm.

We previously showed ATP-stimulated proton transport in a PLV enriched fraction (Miranda et al, 2010, Fig. 6B) supporting activity of the V-H^+^ATPase in the PLV. We tagged the a1 subunit with the Green fluorescent protein mNeon (Sidik et al., 2016) and the parasites expressing VHa1-mNeon-Green were loaded with the lysosomal marker LysoTracker, which labels acidic compartments. LysoTracker co-localized with the VHa1-mNeon at the PLV in extracellular tachyzoites, confirming the acidic nature of the PLV and the co-localization with the V-H^+^-ATPase (**Fig. 1E**).

We wondered if the change in localization of the *a*1 subunit was also observed with other complex subunits. We localized subunits E (TgGT1_305290; henceforth VHE) and G (TgGT1_246560; henceforth VHG) following a similar genetic strategy as the one used for VHa1. Subunits E and G form part of the peripheral V_1_ domain and are involved in establishing the stator of the V-H^+^-ATPase complex and interact with the <u>r</u>egulator of <u>A</u>TPase of <u>v</u>acuoles and <u>e</u>ndosomes (RAVE) complex, which aides in V-H^+^-ATPase assembly (Smardon et al., 2002). Both VHE and VHG subunits localized to the plasma membrane as well as with TgVP1 in both intracellular and extracellular tachyzoites (**Figs. 1F, G, and S2E-F**). In addition, we tagged subunit G (VHG) with a Ty tag in the parasites expressing VHa1-3HA (**Fig. S2D**) to study co-localization. We showed by IFAs almost perfect co-localization of anti-Ty and anti-HA, strengthening the evidence for the interaction between subunits of the complex (**Fig. S2D**). Our results clearly established that the *T. gondii* V-H^+^-ATPase localizes to the plasma membrane and the PLV, an endosomal compartment especially evident in extracellular parasites.

### VHa1 can partially rescue growth in *a* subunit-deficient yeast

The specific function of the VHa1 gene product as part of the proton pumping activity of the V-H^+^-ATPase was investigated by expressing the *Toxoplasma vha1* gene in *Saccharomyces cerevisiae Δvph1Δstv1* mutants (Perzov et al., 2002). These mutants do not express either of the *a* subunits (Vph1p and Stv1p, which localize to the yeast vacuole or Golgi, respectively) and are unable to grow at pH ≥ 7 due to the lack of the V-H^+^-ATPase-dependent acidification of the yeast vacuole (Manolson et al., 1994). The *T. gondii vha1-HA* gene was cloned into the galactose inducible expression plasmid, pYES2/NT C, and transformed into the *Δvph1Δstv1* yeast strain (**Fig. 2A**) to examine the ability of VHa1 to complement the growth phenotype of this strain.

**FIGURE 2:**
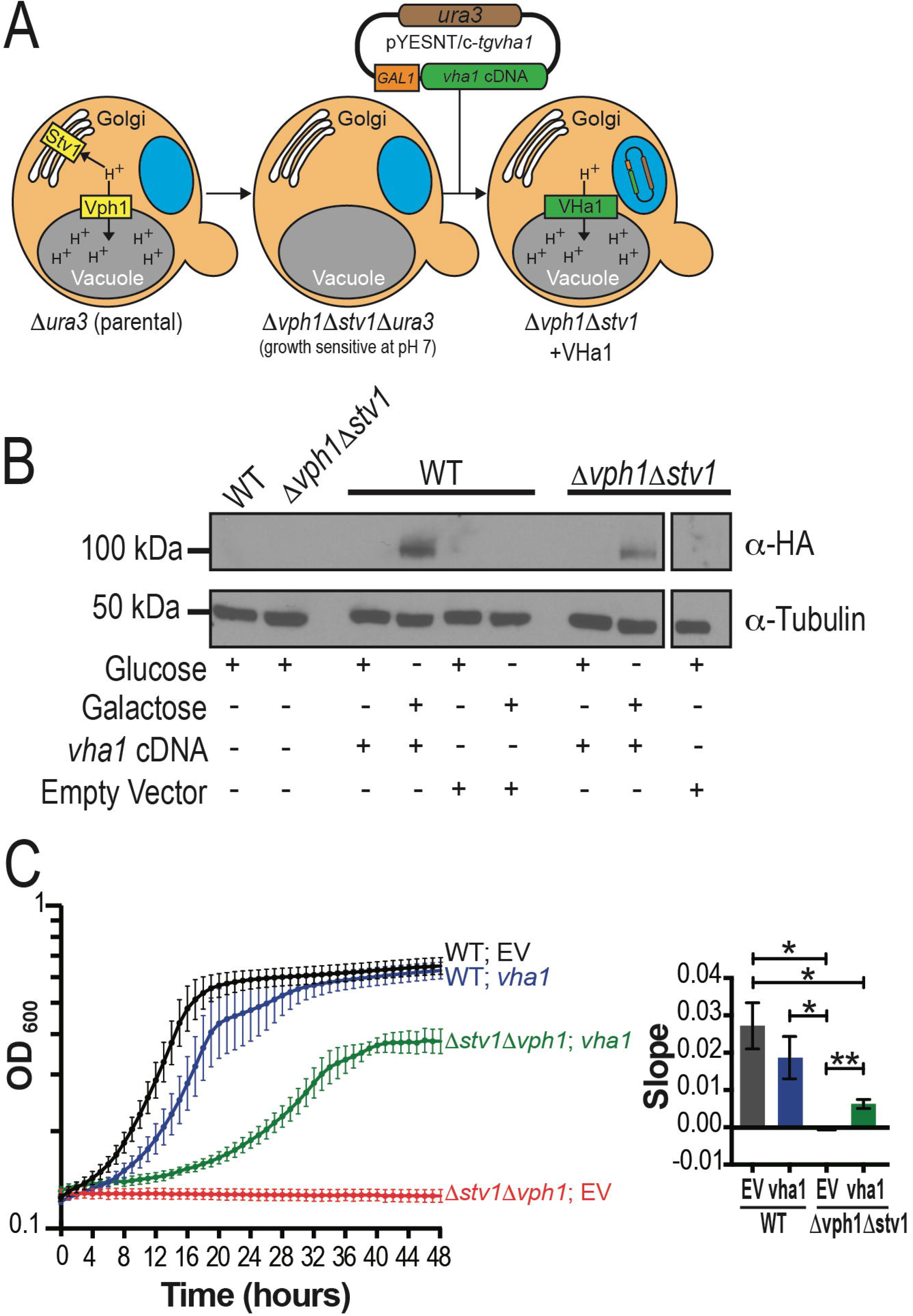
Functional analysis of VHa1 by complementation of mutant *Saccharnmyces cerevisiae*. A) Scheme of the strategy used for complementation of *vph1ΔstvlΔ Saccharomyces cerevisiae* with the *vha1* gene. GAL1, galactose inducible promoter; *ura3*, metabolic marker that contains the *ura3* gene. B) Western blot of *vph1ΔstvlΔ* yeast lysates grown on glucose or galactose (pH 5.5) transfected with a pYES2 vector containing the *vha1-HA* cDNA. Control strains were transformed with an empty pYES2/NT C vector or no vector. C) Growth of WT and *Δvph1Δstvl* yeast harboring *vha1* cDNA (vha1) or empty vector (EV) grown in CSM-ura pH 7.0 with 2% galactose for 48 hours. Data are from three independent trials done in triplicate. Quantification of the slopes during exponential growth (4-16 hours in WT yeast and 8-32 hours in *Δvph1Δstvl* yeast) using one-way ANOVA test where \**P* < 0.05 and \*\**P* < 0.01.

Growing yeast in CSM-ura media at pH 5.5 with 2% galactose resulted in expression of VHa1-HA, while no expression was observed when using 2% glucose (**Fig. 2B**). While the *vph1Δ stv1Δ* strain is incapable of growth on CSM-ura medium at pH 7.0 (**Figs. 2C and S3A**), expression of VHa1-HA partially complemented the growth of the *vph1Δstv1Δ* strain on this medium (**Figs. 2C and S3A**); growth of wild type yeast expressing VHa1-HA was not significantly different from that of a wild type strain harboring the empty vector control (**Figs. 2C and S3A**). Quantification of the slopes during exponential growth (4-16 hours in WT yeast and 8-32 hours in *vph1Δstv1Δ* yeast) showed that *vph1Δstv1Δ* expressing VHa1-HA or WT (empty vector or expressing VHa1-HA) grew significantly better than *vph1Δstv1Δ* yeast harboring the vector control (**Fig. 2C**). Using a similar strategy, the gene of the subunit *a*2 *(VHa2)* was also shown to partially complement the growth phenotype of *vph1Δstv1Δ* yeasts (**Figs. S3A**). We localized VHa1 to the yeast vacuole (and other internal compartments) by expressing GFP-VHa1 in these strains and comparing it to the normal vacuolar localization of GFP-Vph1 (**Fig. S3B**). These yeast complementation results showed that the *T. gondii a1* subunit appears to function as part of the yeast V-H^+^-ATPase complex, thus providing genetic evidence of its conserved biochemical function in acidifying intracellular compartments.

### VHa1 is an essential gene and is important to the lytic cycle of the parasite

The yeast complementation results showed that VHa1 functions as part of the V-H^+^-ATPase multisubunit complex, and we therefore wanted to study the role of the V-H^+^-ATPase in *T. gondii*. We generated a conditional *iΔvha1-HA* mutant parasite cell line by using a tetracycline-regulated transactivator system and the parental cell line TatiΔku80 *(TatiΔku80)*, which combines the high efficiency of homologous recombination (Fox et al., 2009) with regulated gene expression (Sheiner et al., 2011) (**Figs. 3A and S4A**). We confirmed proper insertion and orientation of the promoter by PCR analysis (**Fig. S4B**). VHa1 protein expression was undetectable by Western blot analysis after 24 hours of growth in media containing anhydrotetracyclin (ATc), thus confirming the regulatory function of the promoter on *vha1* (**Fig. 3A**). Subsequent immunofluorescence assays showed that 3 days of ATc treatment resulted in the signal for the VHa1 to be undetectable (**Fig. S4C**, *+ATc)*. Simultaneously, we created a strain of *T. gondii* that constitutively expressed *vha1* from a tubulin promoter at the neutral *uprt* gene locus (Donald and Roos, 1995) in the *iΔvha1-HA* background, to serve as a complementation control, termed *iΔvha1-HA-CM* (**Fig. S4D**); these cells expressed untagged VHa1 even as ATc suppressed the expression of the *vha1-HA locus* (**Fig. 3B**). Note that the immunoblot shown in Fig. 3B was performed using anti-HA (left panel) and the polyclonal antibody anti-VHa1 generated in this work (right panel) (see Materials and Methods) (**Fig. S2H**).

**FIGURE 3:**
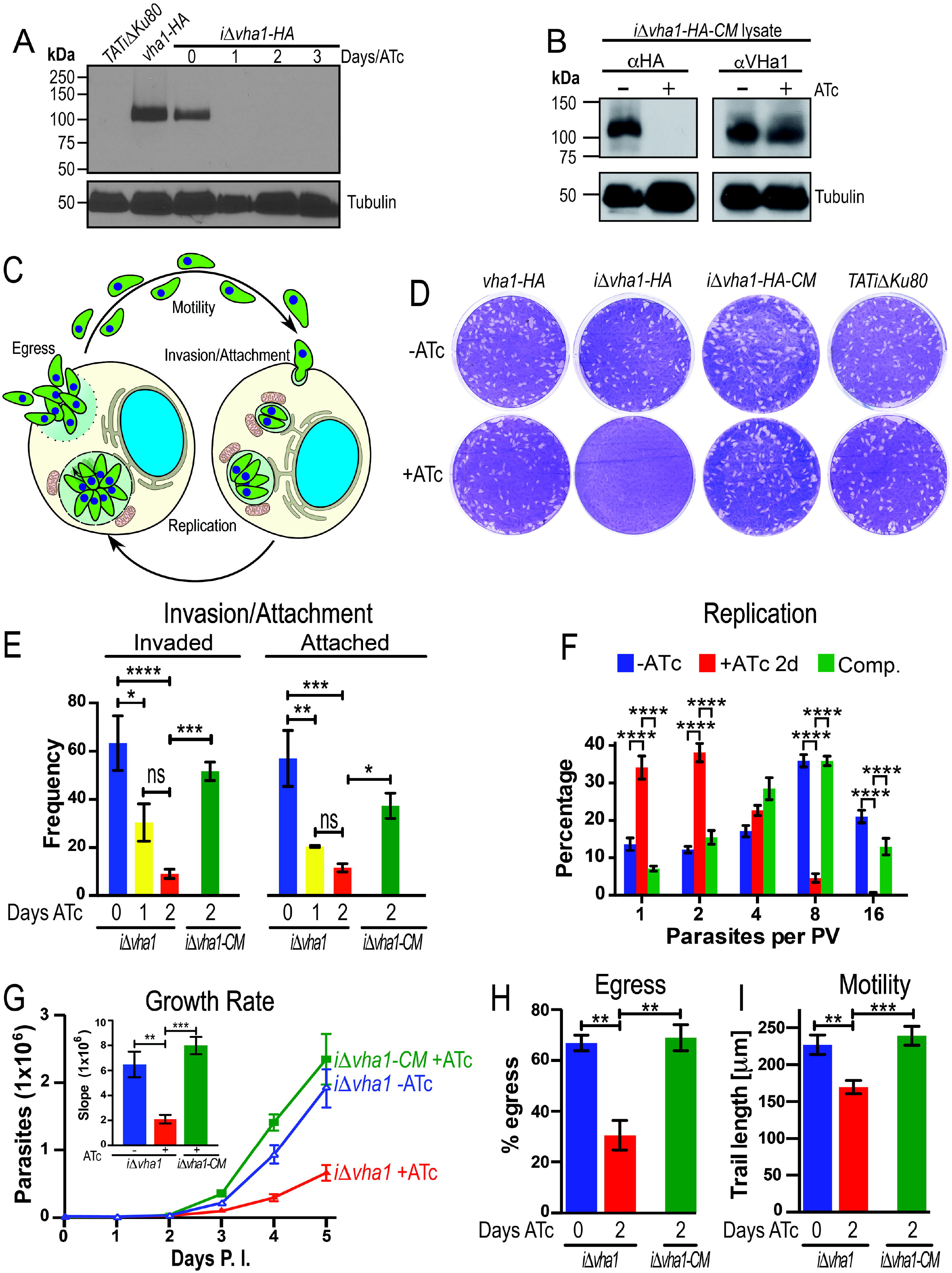
The V-H^+^-ATPase is important to the lytic cycle of *T. gondii*. A) Western blot analysis of parental *(TATiΔku80), vha1-HA*, and *iΔvha1-HA* with or without ATc using rat anti-HA. Anti-tubulin was used as loading control. B) Western blot analysis of lysates of *iΔvha1-HA-CM* parasites showing lack of expression of endogenous VHa1 in the presence of ATc (anti-HA which reacts against the endogenous VHa1) and the expression of the extra copy of VHa1 from the *uprt* locus in the complemented cell line (mouse anti-VHa1 shown in Fig. S2H). C) Overview of the important steps in the lytic cycle. D) Plaque assays of tachyzoites of the HA tagged *(vha1-HA)*, promoter insertion mutant *(iΔvha1-HA)*, complemented *(iΔvha1-HA-CM)*, and parental (TATiΔku80) clones grown in the absence (-ATc) or presence (^+^ATc) of 0.5 μg/ml Anhydrotetracycline (ATc). Each well was infected with 200 parasites and parasites were allowed to form plaques for 8 days at which time they were fixed and stained with crystal violet. E) Red-green assay for quantification of attachment and invasion of *T. gondii* to HFF cells. Data were compiled from 3-4 independent experiments counting ten randomly selected fields per clone. F) Growth kinetics of *iΔvha1-HA* and *iΔvha1-CM* with or without ATc. 115-150 parasitophorous vacuoles were counted per clone in three independent trials with the number of parasites enumerated. G) Approximately 4,000 tdTomato-expressing parasites were grown in confluent hTERT cells in a 96 well plate for 5 days. Data are presented as the average of three independent trials done in triplicate. Inset: slopes from days 2-5 were determined. H) Egress triggered with 10 μM nigericin, which was added to intracellular parasites for 30 min at 37°C. Data were compiled from 3 independent experiments counting ten randomly selected fields per clone and comparing them to a DMSO control. Only parasitophorous vacuoles with 2 or more parasites were enumerated. I) tdTomato-expressing parasites were resuspended in ringer buffer without Ca^2+^ and motility was stimulated by adding 1.8 mM Ca^2+^. The average length (in microns) parasites traveled is reported from 5-6 individual parasites from 3 independent trails. Values are means ± SEM and data were analyzed by GraphPad Prism 6. E) and F) were compared with two-way ANOVA test; G), H), and I) were compared with one-way ANOVA test. \**P* < 0.05; \*\**P* < 0.01; \*\*\**P* < 0.001, \*\*\*\**P* < 0.0001.

The pathogenicity of *T. gondii* stems from its lytic cycle, which is initiated by active invasion of a host cell, subsequent replication inside a parasitophorous vacuole, and egress (**Fig. 3C**). To establish whether VHa1 is important for parasite growth, we first compared the growth of *vha1-HA, iΔvha1-HA, iΔvha1-HA-CM*, and the parental strain TATiΔku80 in parallel plaque assays (**Fig. 3D**). Only *iΔvha1-HA* showed a clear defect in the formation of plaques when grown in the presence of ATc, which suppresses the expression of *vha1*, demonstrating its critical role in at least one major step of the lytic cycle. *iΔvha1-HA* parasites pre-incubated with ATc for 3 or 4 days, before washing away the ATc, were unable to complete their lytic cycle, although some growth remained after a 2 day treatment (**Fig. S5A**, *green). iΔvha1-HA* parasites grown with ATc for three days were still viable as detected by trypan blue exclusion (**Fig. S5B**). To ensure that the parasites incubated with ATc were viable, we tested trypan blue exclusion of the *iΔvha1-HA* parasites grown with ATc for 0 or 3 days and according to the result shown in Fig. S5B they were almost 100% viable. Based on these results, we studied the impact of down-regulating V-H^+^-ATPase on a variety of biological phenotypic features with 1, 2, or 3 days (but no longer than 4 days) after ATc treatment. We reasoned that longer exposure of ATc than 4 days would likely result in stronger phenotypic characteristics, but could be the result of nonspecific effects.

We next analyzed each step of the lytic cycle and found that 2 days treatment with ATc was sufficient to reduce invasion of host cells by 7-fold and attachment by 5-fold (**Fig. 3E**); these defects were ameliorated in the *iΔvha1-HA-CM* cell line. By introducing cytosolic tdTomato into *iΔvha1-HA and iΔvha1-HA-CM*, we could analyze growth kinetics of *T. gondii* cellular replication during infection. Parasites were pre-incubated (+ATc) or not (-ATc) with ATc for 24 hours and passaged to new fibroblast cells with or without ATc, respectively. After 24 h of exposure to ATc, followed by another 24 h in a new passage (48 h of total ATc treatment in ^+^ATc conditions), there was a significant reduction in the ability of *iΔvha1-HA +*ATc clones to replicate, when compared to the *iΔvha1-HA-CM* line (**Fig. 3F**). Furthermore, *iΔvha1-HA* parasites grew much more slowly in the presence of ATc, in contrast to either *iΔvha1-HA-CM* in the presence of ATc, or *iΔvha1-HA* cells in the absence of ATc treatment (**Figs. 3G and S5C**). We studied egress by stimulating intracellular tachyzoites with nigericin or saponin and calcium (Borges-Pereira et al., 2015) and the *iΔvha1-HA+ATc* cells showed a significant delay in their response to these inducers of egress (**Figs. 3H** and **S5D**). There was a significant difference in the % of lysed parasitophorous vacuoles counted after 30 min of exposing intracellular parasites to 10 μM nigericin (66.9% and 68.9% for the *iΔvha1-HA-ATc* and the *iΔvha1-HA-CM^+^ATc*, respectively, and 30.5% in *iΔvha1-HA+ATc)* (**Fig. 3H**). We also measured time to egress after induction with 2 mM Ca^2+^ in the presence of 0.01% saponin to permeabilize host cells. *iΔvha1-HA-ATc* and *iΔvha1-HA-CM+ATc* exited host cells at 303 ± 35 and 342 ± 80 seconds, respectively, while *iΔvha1-HA+ATc* took an average of 527 ± 45 seconds (**Fig. S5D**). We also evaluated motility, an essential part of the lytic cycle of *Toxoplasma* and found that when knocking down the expression of the *vha1* gene with ATc, *(iΔvha1-HA+ATc)*, the parasites are not able to travel as far as the wild type ones (**Fig. 3I**). These mutants traveled 57 μm less than the -ATc parental and 69 μm less than the complemented strains incubated with ATc (Fig. 3I). In summary, these results show that the activity of the V-H^+^-ATPase impacts every major step of the lytic cycle of *Toxoplasma*.

### The role of VHa1 in monitoring intracellular pH

Intracellular pH (pH_i_) must be strictly controlled because of the narrow optimum pH of most intracellular enzymatic processes (Demaurex, 2002). Because of this, cells regulate their pH_i_ by active transport of H^+^ across membranes. Our lab previously showed that the maintenance of the cytoplasmic pH of *Toxoplasma* tachyzoites was sensitive to bafilomycin A_1_, a specific inhibitor of the V-H^+^-ATPase (Moreno et al., 1998) and we proposed a role for the V-H^+^-ATPase in pH_i_ regulation. We therefore tested pH_i_ in the *iΔvha1* mutants grown in the presence of ATc. To determine the role that the V-H^+^-ATPase plays in pH_i_ regulation, we used the chemical pH indicator BCECF and its acetoxymethyl ester BCECF-AM to measure external and internal changes in pH, respectively (**Figs. 4A, B**). We measured intracellular pH of extracellular tachyzoites under several extracellular pHs and we did not find a significant difference in the pHi values of *iΔVHa1+ATc* compared to the parental and complemented lines (**Fig. 4B**). Note that the changes in extracellular pHs to which the parasites are exposed vary between 5.5 and 8.5. This experiment shows that these parasites are still able to regulate pH under these extracellular conditions, which is reasonable considering the importance of maintaining a physiological pH_i_ level. We next looked at extrusion of protons from intact cells (Fig. 4C). This is performed by using the free acid form of BCECF in a weakly buffered solution (Pace et al., 2011). This protocol allows to measure changes in extracellular pH (pH_e_), that results from proton extruded from the cell into the extracellular milieu. Addition of glucose, which stimulates the glycolytic activity of tachyzoites, resulted in medium acidification (**Fig. 4C** *note the increase in slope of green and blue tracings)*. Changes in pH_e_ of −0.40 to −0.46 pH units were observed with *iΔvha1-* ATc or with iΔvha1-CM^+^ATc tachyzoites. However, an average change in pH_e_ of only −0.19 pH units was obtained in *iΔvha1+ATc* tachyzoites (**Fig. 4C and inset**). The proton extrusion activity was completely blocked by bafilomycin A_1_, and the change in extracellular pH was not significantly different between *iΔVHa1*-ATc, *iΔVHa1+*ATc, and *iΔVHa1-CM+*ATc parasites under these conditions (**Fig. 4D**). These data indicate an important role for the *T. gondii* V-H^+^-ATPase-dependent in the extrusion of protons upon stimulation of metabolic activities with glucose.

**FIGURE 4:**
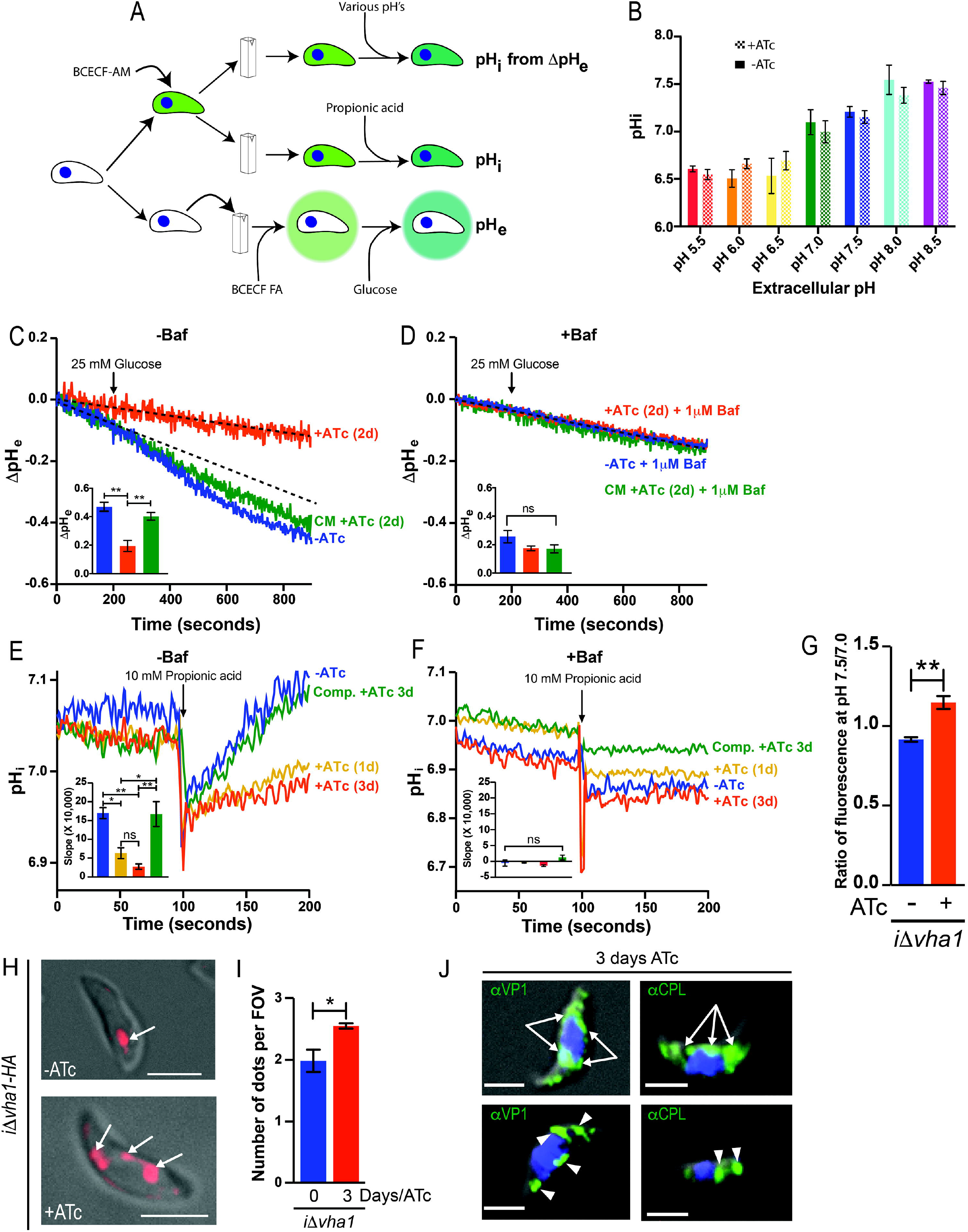
Function of VHa1 in *T. gondii* tachyzoites. A) Scheme of the protocol used to test the effects on intracellular pH (pHi), the effect of changing extracellular pH on internal pH (pH_i_ from ΔpH_e_), and proton extrusion (pH_e_). B) Intracellular pH measurements of *iΔvha1-HA* cells at various extracellular pH’s. Parasites were loaded with the pH indicator BCECF-AM and fluorescence measurements were done as described in Materials and Methods. Data from from 3 independent trials. C) Representative tracings of proton extrusion of *iΔvha1-HA* with or without ATc with the addition of 5 mM glucose were obtained using BCECF free acid in a weakly buffered solution. Changes in pH were followed as described in Materials and Methods. Results are from 3-6 independent trials. Dashed line represents the slope of -ATc before the addition of glucose. D) Same as C but with cells pretreated with 1 μM bafilomycin for 3 min. C and D insets show the quantification of slope changes between 0-900 sec. E) Acid load and pH recovery of parasites loaded with BCECF-AM. 10 mM propionic acid were added where indicated and acidification was followed by pH recovery. Parasites tested were *iΔvha1-HA* grown in the presence of ATc for 0, 1, 3 days and *iΔvha1-HA-CM* (3 days ATc). Results are from 3-4 independent trials. F) Same as E but cells were pretreated with 1 μM bafilomycin for 3 min. E and F insets show the quantification of the slope from 105 to 200 sec from 3 independent trials. G) Membrane potential measurements of *iΔvha1-HA* cells grown with or without ATc for 2 days and incubated with bisoxanol. The fluorescence measurements at pH 7.5 to 7.0 were compared and are from 3 independent trials. H) *iΔVHa1* parasites incubated with or without ATc (0 or 3 days, respectively) and loaded with 10 μM LysoTracker red for 30 min at 37°C in BAG I) Quantification of compartments stained by LysoTracker. 5-15 cells per field of view were counted and a minimum of 8 fields considered. Graph shows the quantification of puncta counted in *iΔVHa1* parasites in the presence and absence of ATc. J) IFA of *iΔVHa1* parasites incubated with ATc for 3 days and probed with anti-VP1 or anti-CPL. Arrows point to endoplasmic reticulum localization and arrowheads point to discrete puncta of VP1 or CPL staining. Statistical analysis was performed using a student’s T-Test (G, I) or one-way ANOVA (B, C-F) where \**P* < 0.05; \*\**P* < 0.01; \*\*\**P* < 0.001, \*\*\*\**P* < 0.0001, ns, not significant.

Exposing *iΔVHa1-ATc* tachyzoites to 10 mM propionic acid causes transient acidification of the cytosol, which is quickly recovered to normal values within 50 s of the acid pulse (**Fig. 4E**, *blue tracing*). The recovery of the intracellular pH of *iΔVHa1*+ATc parasites after acid treatment was very slow and 100 seconds post-treatment their intracellular pH was still very low, while control parasites were completely recovered (**Fig. 4E**, *yellow and red tracings*). *iΔVHa1-CM* parasites were able to restore its pH_i_, almost as fast and efficiently as the parental parasites even in the presence of ATc (**Fig. 4E**, *green tracing*). Furthermore, bafilomycin A_1_ blocked recovery of pHi in all cell lines (with and without ATc and complemented strain) (**Fig. 4F**), showing that the V-H^+^-ATPase functions to protect *T. gondii* from acid stress (and likely from normal intracellular acid production) by pumping protons out of the parasite cytosol.

Deficient proton pumping at the plasma membrane may also impact the membrane potential of tachyzoites. We used the membrane potential-sensitive fluorescent probe bisoxonol (Moreno et al., 1998). The *iΔVHa1-HA*+ATc were modestly depolarized (higher fluorescence ratio) when compared with parasites grown without ATc (**Fig. 4G**). These data indicated that the proton pumping activity of the V-H^+^-ATPase generated a H^+^ gradient at the plasma membrane that contributes partially to the build-up of the membrane potential. It is likely that other mechanisms are at play and prevent cells from becoming completely depolarized. These results underscore the functional activity of the V-H^+^-ATPase at the plasma membrane by extruding protons upon cytoplasmic acidification.

Lysotracker, a fluorescent indicator of acidic compartments co-localized with VHa1 in the PLV as shown in Fig. 1E, supports the acidic nature of the PLV as well as the potential role of the V-H^+^-ATPase in its acidification. We next tested Lysotracker in the *iΔVHa1*-HA+ATc, which showed an intriguing phenotype (**Fig. 4H, I**). *iΔVHa1*-HA+ATc parasites showed higher number of vacuoles labeled with the fluorescent indicator. We interpreted this result as a defect in the biogenesis of the PLV so we next performed IFAs of *iΔVHa1*-HA+ATc parasites with the PLV markers VP1 and CPL. Mutant parasites showed labeling of VP1 and CPL in vesicles distributed all through the cell (**Fig. 4J *arrowheads***) and they even labeled the endoplasmic reticulum (**Fig. 4J *arrows***). Our interpretation is that the PLV is not forming properly in extracellular *iΔVHa1*-HA+ATc tachyzoites, probably due to defective vesicle fusion (**Fig. 1 iii and iv**). The vesicular labeling observed by VP1 staining was similar to the one observed with LysoTracker, supporting that the defective formation of the PLV was likely leading to an accumulation of VP1 vesicles, which were acidic but failed to fuse to form the PLV. We were not able to evaluate the difference in acidification because of the non-quantitative nature of LysoTracker. To quantify a change in the pH of VHa1 depleted parasites, we sent a genetically encoded pH sensing protein indicator (mCherry-SEpHluorin) to the PLV (see supplemental movie 1). However, because the fragmented nature of the PLV in extracellular parasites, it was not possible to determine pH changes in the resulting vesicles (data not shown). Our data supports a role for the V-H^+^-ATPase in trafficking of vesicles to or from the PLV.

### VHa1 is important for the localization, maturation, and secretion of micronemes

Micronemes are secretory organelles involved in the invasion of host cells by tachyzoites (Carruthers et al., 1999). Several microneme proteins undergo proteolytic maturation while traveling to the micronemes and a role for endosomal compartments in this maturation has been previously demonstrated (Harper et al., 2006). The V-H^+^-ATPase translocates from the plasma membrane to an internal large and dynamic vacuole, the PLV, in extracellular tachyzoites. The PLV was previously shown to have lytic characteristics and harbors microneme maturase activity (Miranda et al., 2010; Parussini et al., 2010). Parasite attachment and invasion is associated with the secretion of micronemes (Carruthers et al., 1999), and since *iΔVHa1* parasites display defects in attachment and invasion, we examined the role of the V-H^+^-ATPase on microneme secretion. We tested the release of the microneme protein, MIC2, into the excreted/secreted antigen fraction collected from supernatants of extracellular tachyzoites without stimulation (**Fig. 5A**, *constitutive*) or after induction of secretion by ethanol (**Fig. 5A**, *1% ethanol). iΔVHa1* (+ATc) parasites were not able to secrete micronemes constitutively and they responded poorly to stimulation by ethanol (**Fig. 5A**). The amount of total MIC2 protein in the *iΔvha1* parasite lysate was similar to those of the parental and complemented strains. **Fig. 5B** shows the quantification of three independent experiments using the constitutively secreted GRA1 protein as control. We found that after 3 days of incubation with ATc the *iΔvha1* tachyzoites are deficient at secreting MIC2 while secreting normal levels of GRA1 (indicating that parasites are viable) (**Fig. 5B**). Complementation of the *iΔvha1* cell line with exogenous *vha1* (*iΔvha1–CM*) restored normal levels of MIC2 secretion even when grown for 3 days in the presence of ATc. We also tested the secretion of the apical membrane antigen 1 (TgAMA1), a transmembrane protein that localizes to the parasite’s micronemes (Hehl et al., 2000). Secretion of TgAMA1 was also significantly reduced in the *iΔvha1+ATc* line (**Fig. S6**).

**FIGURE 5:**
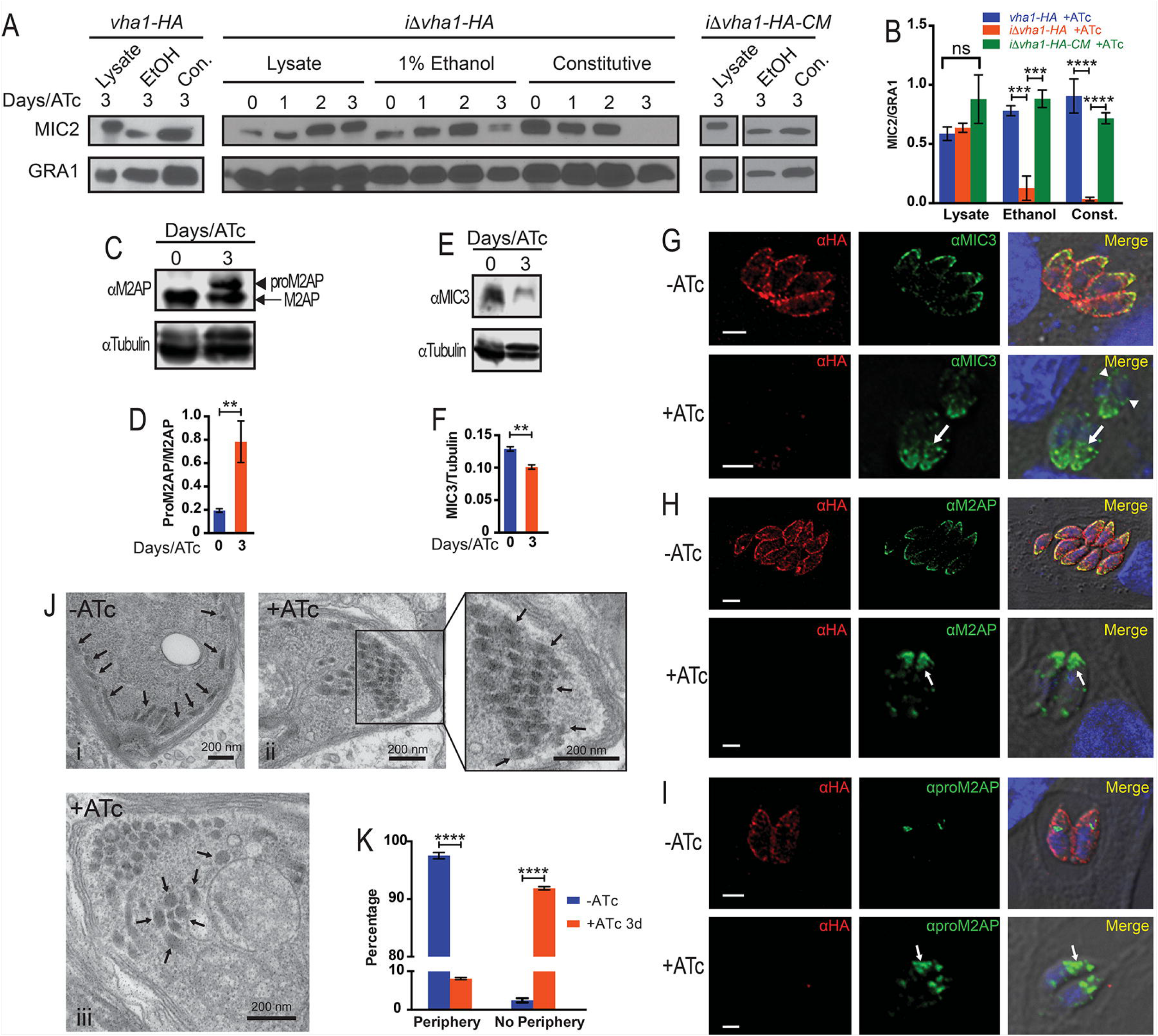
V-H^+^-ATPase activity and microneme secretion. A) Western blots showing microneme secretion experiments of *vha1-HA, iΔvha1-HA*, and *iΔvha1-HA-CM* parasites. Parasites were grown for the indicated time with ATc, harvested, and washed. Parasites were incubated in invasion media alone for constitutive expression (cons) or in the presence of 1% ethanol (EtOH). Lysate samples were also included. Aliquots of the supernatants were analyzed by Western blots with anti-MIC2 or anti-GRA1. B) Quantification of MIC2 secretion by pixel density from ImageJ (ratio of MIC2/GRA1) from 4-5 independent trials. Values are means ± SEM. C) Western blots of total lysates of *iΔvha1-HA* parasites grown with and without ATc showing the increase in the signal for proM2AP (arrow head) compared to mature M2AP (arrow). D) Ratio of the intensity of the bands corresponding to proM2AP and M2AP. Bar graphs shows the quantification of at least three independent experiments. E) Western blots of total lysates of *iΔvha1-HA* parasites grown with ATc for 3 days or without ATc (0) probed with anti-MIC3. F) Quantification of the ratio of the intensity of the bands for MIC3/tubulin from three independent experiments. G) IFAs with anti-MIC3 and anti-HA showing localization of MIC3 in *Δvha1-HA* parasites grown with or without ATc. Arrowheads point toward perinuclear labeling and arrows point staining not at the periphery. H) IFAs with anti-M2AP and anti-HA showing the localization of M2AP in *iΔvha1-HA* parasites grown with and without ATc. I) IFAs with anti-proM2AP and anti-HA in *iΔvha1-HA* parasites grown with or without ATc. J) Transmission Electron microscopy of *iΔvha1-HA* parasites grown without ATc (i) or in the presence of ATc for 3 days (ii-iii). i) Arrows point toward peripheral micronemes and at the apical end of the parasite. ii) Micronemes are indicated by the arrows toward the apical end in parasites grown with ATc for 3 days. iii) micronemes are more centrally located in parasites incubated with ATc for 3 days. K) Quantification of EM images from three independent preparations comparing the localization of micronemes of *iΔvha1-HA* grown in the presence or absence of ATc for three days. Only 1 periphery microneme 2 μm from apical tip was considered to confirm a positive localization at the periphery. Approximately 59-72 parasites were enumerated from 3 independent trials. F, H were quantified by pixel density using Image Studio from 3 independent trials. Graph error bars are SEM. Statistical analysis were performed using a one-way ANOVA where \*\**P* < 0.01; \*\*\**P* < 0.001, \*\*\*\**P* < 0.0001, ns, not significant.

MIC2 forms an heterohexameric complex with M2AP (MIC2-associated protein) and the complex MIC2-M2AP plays fundamental roles in gliding motility and invasion (Harper et al., 2006). Both proteins are proteolytically processed during their transport to the micronemes (El Hajj et al., 2008; Garcia-Réguet et al., 2000). M2AP is initially translated with a propeptide that is removed in an intracellular endosomal compartment and blocking the removal of the propeptide would affect trafficking of the M2AP-MIC2 complex to the periphery. With the aim to study the impact of proton pumping on the processing and trafficking of M2AP we investigated its maturation in the *iΔvha1* (+ATc) cells treated with ATc for 3 days and we observed a significant accumulation of the immature form (**Fig. 5C-D**). Controls with the parental line *vha1-HA*, showed no defects in the maturation of M2AP when grown with or without ATc (**Fig. S7A, B**). MIC3 is an homodimeric adhesin that is synthesized as a precursor that is proteolytically processed during its traffic through the secretory pathway on its way to the micronemes. This processing is important for the expression of the binding function of the protein (Cérède et al., 2002). Mature MIC3 abundance was significantly reduced in *iΔvha1* (+ATc) when compared to controls without ATc (**Figs. 5E, F**) or to the parental line with ATc (**Fig. S7A, C**).

We next looked at the distribution pattern of microneme proteins by IFA in the *iΔvha1-HA+ATc* mutants (**Figs. 5G-I**). There was a notable difference in the distribution of MIC3 labeling in the *iΔvha1* (+ATc) parasites (**Fig. 5G**). The increased level of MIC3 in perinuclear areas and their accumulation toward the posterior end in vesicles likely destined to the secretion pathway was clearly evident in the *iΔvha1*+ATc parasites (**Fig. 5G**, *arrows*). This was distinct from the neatly distributed labeling of MIC3 at the periphery of parasites that express normal levels of VHa1 (*iΔvha1*-ATc) (**Fig. 5G**). It was previously reported that unprocessed MIC3 builds up in perinuclear compartments such as the ER and/or Golgi compartments and accumulates in the secretory pathway during parasite division (El Hajj et al., 2008). These data point towards retention of MIC3 in the ER and secretory pathway because of defective processing without the acidification function of the V-H^+^-ATPase (**Fig. 5G**, *arrowheads*). MIC3 and M2AP are synthesized almost simultaneously but pro-M2AP was mainly observed in endosome-like compartments while pro-MIC3 was observed in both the ER-Golgi and endosome-like compartments (El Hajj et al., 2008). We studied the localization of M2AP in the *iΔvha1*+ATc parasites (**Fig. 5H**) and we also saw a remarkable difference in its distribution. In *iΔvha1*+ATc parasites M2AP was clustered within the central region of the extreme apical end, while *iΔvha1*-ATc showed the typical distribution of labeling at the periphery of the parasite toward the apical end (**Fig. 5H**, *arrows*) similar to the localization of MIC2. Disruption of M2AP (Harper et al., 2006) results in secretory retention of TgMIC2, leading to reduced TgMIC2 secretion from the micronemes and impaired invasion. The localization of MIC2 was also altered and the signal showed an accumulation of vesicles with intense labeling toward the apical end of the cell close to the plasma membrane (**Fig. S8**).

M2AP is initially translated with a propeptide that is removed in an intracellular endosomal compartment and blocking the removal of the propeptide would affect trafficking of the M2AP-MIC2 complex to the periphery. The *iΔvha1* (+ATc) cells treated with ATc for 24 hrs showed a maturation delay of the proM2AP (**Fig. 5C**) and a significant accumulation was detected after 3 days of ATc incubation (**Figs. 5I**). Our data showed that the V-H^+^-ATPase likely acidifies a compartment that is in the path of the maturation pathway of M2AP and MIC3, two microneme proteins that are processed before storage in the microneme compartment (Soldati et al., 2001).

Transmission electron microscopy of *iΔvha1-HA* parasites treated with ATc showed that their micronemes are not able to dock or fuse at the periphery, instead presenting a more central location in the apical end of the tachyzoite (**Fig. 5J**). These micronemes also exhibit a rounded morphology rather than the typical cigar shape (Paredes-Santos et al., 2012; Tomavo et al., 2013) (**Fig. 5J**). We found significantly fewer micronemes at the periphery in the *iΔvha1* (+ATc) cells (**Fig. 5K**). Therefore, it is likely that the V-H^+^-ATPase plays a significant role in microneme maturation, function, and possibly organelle distribution.

### VHa1 is important for the maturation of rhoptry organelles and rhoptry proteins

The majority of characterized rhoptry bulb proteins are initially translated with an ER signal peptide and a pro-peptide that acts as a trafficking signal for the rhoptries (Hajagos et al., 2012; Soldati et al., 1998). Because pro-peptides are proteolytically cleaved within endosomal compartments (Ngo et al., 2004), we next questioned if the deficient activity of the V-H^+^-ATPase would impact the maturation of rhoptries as well. Immunoblots of lysates from *iΔvha1*-ATc and *iΔvha1*+ATc tachyzoites revealed that the maturation of the rhoptry proteins ROP4, ROP7, and TgCA_RP (Chasen et al., 2017) were all disrupted, as demonstrated by a significant increase in the percentage of immature protein (**Figs. 6A, B**). Controls with the *vha1-HA* parental line and the effect of ATc on the maturation of ROP4, ROP7, and TgCA_RP are shown in **Fig. S7D, E**.

**FIGURE 6:**
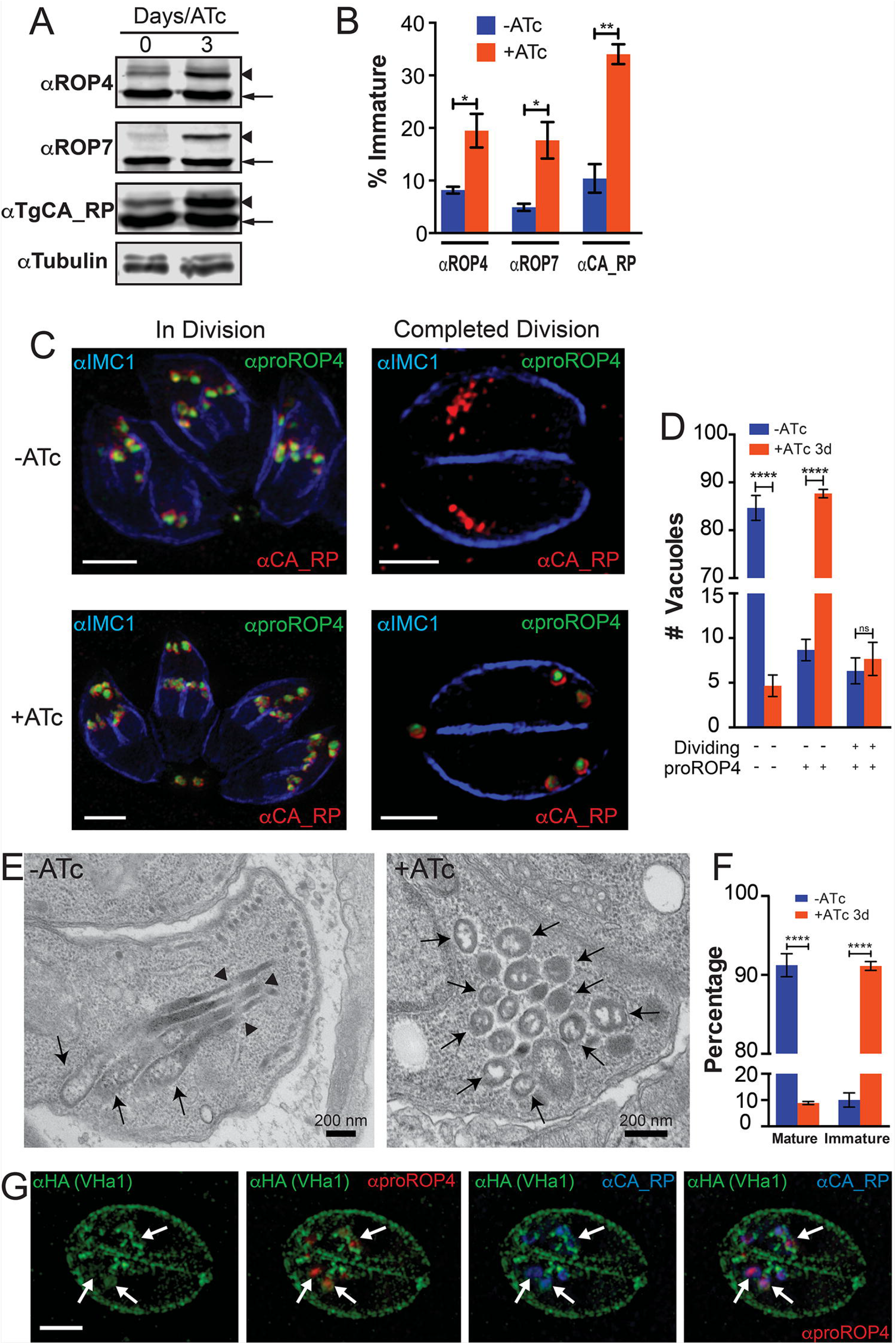
The V-H^+^-ATPase and rhoptry maturation. A) Western blots of total lysates of *iΔvha1-HA* tachyzoites grown with or without ATc (3 days) and probed for the rhoptry bulb proteins ROP4, ROP7, or TgCA_RP. There is an increase in the immature (arrowheads) compared to the mature form (arrows). B) Quantification of percentage of immature ROP4, ROP7, or TgCA_RP showed significant differences when comparing lysates from parasites grown in the presence or absence of ATc. C) Immunofluorescence assays of *iΔvha1-HA* tachyzoites grown with or without ATc. Super-resolution images show that rhoptry protein localization to the mature rhoptries is not disrupted in *dividing* tachyzoites, but rhoptries and rhoptry proteins are not matured properly in tachyzoites that are *not dividing*. Scale bars: 2 μm. D) Quantification of parasitophorous vacuoles with parasites expressing immature and mature rhoptries in *iΔvha1-HA* tachyzoite vacuoles grown with and without ATc. There was a significant increase in the labeling of pro-Rop4 (immature ROP marker) in non-dividing parasites in the presence of ATc (red *columns)*. IMC1 labeling was used to differentiate dividing from non-dividing tachyzoites. E) Routine electron microscopy of a *iΔvha1-HA* tachyzoite grown without ATc (left, *-ATc)* shows normal rhoptries and their characteristic rhoptry neck *(arrowheads)* and bulb regions *(arrows)*. In *iΔvha1-HA* grown in ATc for 3 days, there is a marked absence of mature rhoptries, and accumulation of vesicular immature rhoptry structures *(arrows)*. F) Quantification of the electron microscopy images showing a significant reduction in the number of tachyzoites containing mature rhoptries after treatment with ATc. Approximately 52-73 parasites were enumerated from 3 independent trials. G) Super resolution IFA showing colocalization of VHa1 with proROP4 (1:500) and TgCA_RP (1:1000) in intracellular parasites. Statistical analysis were performed using a Student’s T-test where \**P* < 0.05; \*\**P* < 0.01; \*\*\*\**P* < 0.0001, n. s., not significant.

We next performed immunofluorescence assays of *iΔvha1-ATc* and *iΔvha1+*ATc tachyzoites with antibodies against the pro-peptide of ROP4 (α-proROP4), which labels immature rhoptries (Sakura et al., 2016). We found a significant increase in the number of *iΔvha1* tachyzoites that contained immature rhoptries after exposure to ATc (*iΔvha1*+ATc). This was notable, because the majority of these tachyzoites were not in the division stage of the cell cycle, as determined by IMC labeling (**Fig. 6C**, *not dividing*). In dividing tachyzoites, the rhoptry morphology was indistinguishable between *iΔvha1*-ATc and *iΔvha1*+ATc tachyzoites, however there was an increase in what appeared to be intact nascent rhoptries in the residual body of *iΔvha1*+ATc vacuoles (**Fig. 6C**, *dividing*). Trafficking from the ER appeared to be normal as indicated by the localization of rhoptry bulb proteins to the immature rhoptries. In order to quantify the rhoptry maturation defect, we counted vacuoles containing tachyzoites labeled with the immature rhoptry marker, α-proROP4. We observed a significant increase (>80%) in *iΔvha1* parasite vacuoles labeled by α-proROP4 after 3 days of ATc treatment (**Fig. 6D**). It is notable that the number of vacuoles containing tachyzoites with developing daughter cells (labeled with α-IMC1) was not significantly different in *iΔvha1*+ATc mutants; suggesting that this step of endodyogeny is not affected by the disrupted activity of the V-H^+^-ATPase. Routine electron microscopy of *iΔvha1*+ATc parasites showed a striking phenotype where there is no evidence of mature rhoptries (**Figs. 6E, F**). This data showed that the diminished activity of the V-H^+^-ATPase negatively impacted the maturation of rhoptry proteins and prevented the morphological changes associated with the formation of the characteristic elongated mature rhoptries.

We next performed co-localization studies with α-proROP4 (for labeling immature rhoptries), α-CA_RP (to label mature rhoptries) and α-HA in *iΔvha1-HA* parasites and looked at the images using super resolution microscopy. We observed intracellular labeling of VHa1 to the periphery of vesicles surrounding immature rhoptries (**Figs. 6G**, *arrows* **and S9A**). VHa1 only associated with immature rhoptries but did not co-localize with mature rhoptries (**Fig. S9B**). With the aim to confirm that the whole V-H^+^-ATPase complex localizes to the immature rhoptries, we performed co-localization with *α*-proROP4 and anti-HA of parasites that had the genes for the E and G subunits HA-tagged. We found the labeling of E and G also encircled immature rhoptries, indicating that the whole complex is present at the immature rhoptries and is likely functional (**Figs. S9C, D**). Immature rhoptries were previously shown to be acidic (pH 3.5-5.5) (Shaw et al., 1998) and our data proposes a mechanism by which they are acidified. It remains to be determined how the V-H^+^-ATPase-labeled vesicles split allowing the mature rhoptry proteins to traffic to the rhoptries while the V-H^+^-ATPase follows a different path to other endosomal compartments.

### The V-H^+^-ATPase and proteolytic activities

We investigated if the activity of the V-H^+^-ATPase would be important for the proteolytic activities present in the PLV or ELCs. Cathepsin L (TgCPL), an enzyme present in the PLV is also proteolytically processed and acidic pH is required for both its maturation and its function (Parussini et al., 2010). It has also been reported to be responsible for the maturational M2AP and MIC3 (Parussini et al., 2010). We looked at TgCPL maturation in *iΔvha1* cells and found that 48 hours or more of ATc treatment was sufficient to increase the accumulation of the immature form of TgCPL (**Figs. 7A, B**). The addition of ATc had no effect on *vha1-HA* parental parasites (**Fig. S7F, G**), suggesting that the maturation defect of CPL is related to the activity of the V-H^+^-ATPase.

**FIGURE 7:**
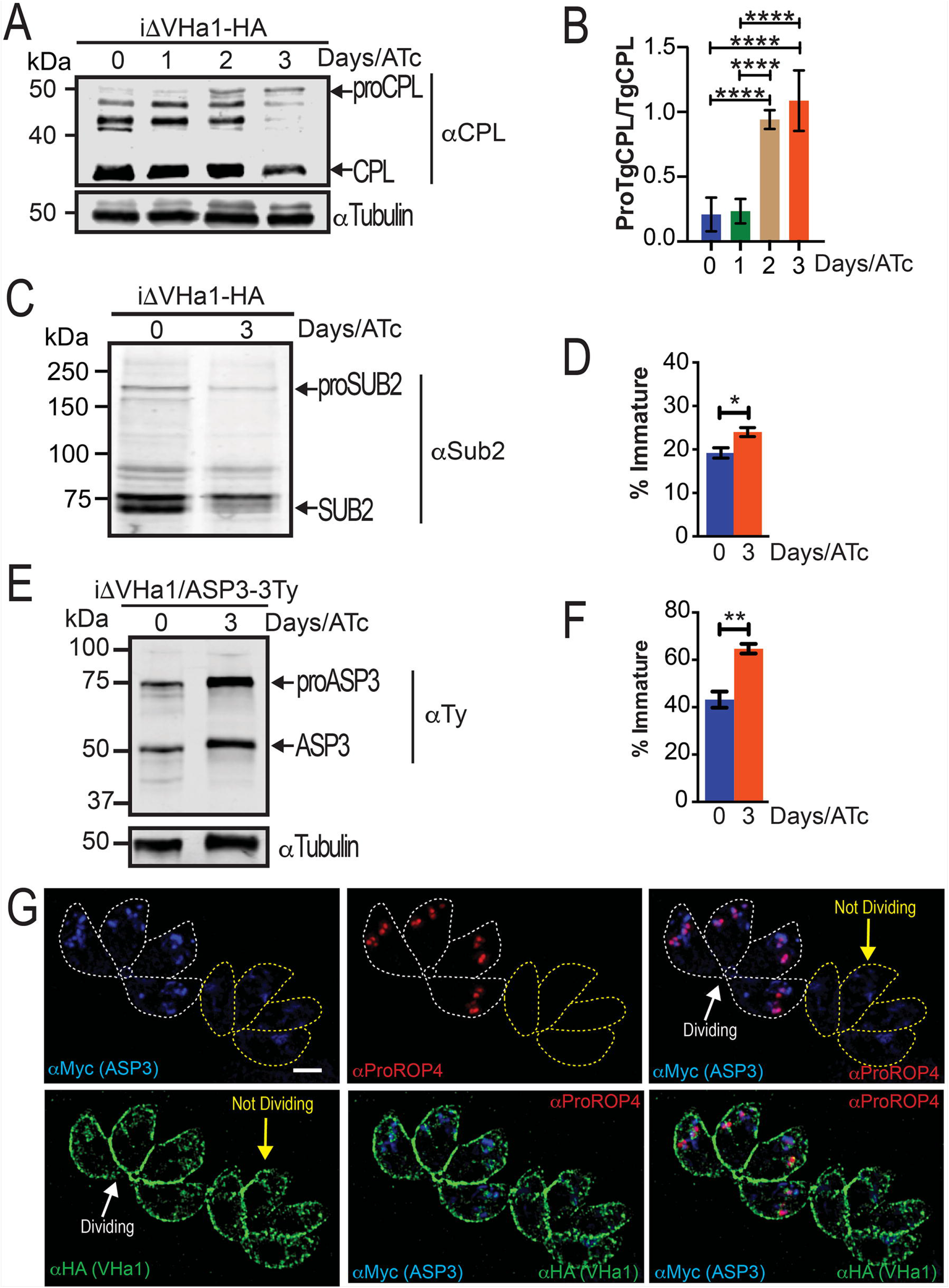
The V-H^+^-ATPase and the maturation of proteases. A) Western blot of *iΔvha1-HA* grown with ATc for the indicated times showing the accumulation of the pro form of CPL compared to the mature form. Molecular weights are in kDa. B) Quantification of the bands shown in A from 4-5 independent trials. Experiments were standardized to tubulin and a ratio of proCPL/CPL was determined by ImageJ. C) Western blots of total lysates of *iΔvha1-HA* tachyzoites grown with or without ATc (3 days) and probed with anti-TgSUB2. D) Quantification of the maturation of SUB2 as percentage of immature form from the total TgSub2. This quantification was done for lysates of *iΔvha1-HA* grown + and – ATc. E) Western blots probed anti-Ty of *iΔvha1-HA* parasite lysates that also express the TgASP3-Ty from the UPRT locus. Clonal parasites were grown with ATc for 0 or 3 days. F) Quantification of the maturation of ASP3 as percentage of the immature form from total ASP3. There was a significant difference in the comparison between lysates from parasites grown in the presence or absence of ATc. G) Super-resolution images show that ASP3 co-localizes with immature rhoptry marker proROP4 and the V-H^+^-ATPase surrounds ASP3 and proROP4 (as indicated with VHa1-HA) in dividing cells (lines outline parasites). Statistical analysis was performed using a one-way ANOVA or student’s T-Test where \**P* < 0.05; \*\**P* < 0.01; \*\*\*\**P* < 0.0001, ns, not significant.

A rhoptry localized subtilisin-like serine protease, TgSUB2 had been proposed to be involved in the maturation of ROP proteins (Miller et al., 2003), although TgSUB2 knock-outs appeared to not show a ROP maturation defect (Dogga et al., 2017). It has been reported that some subtilisin proteases require pH below 7 for their own maturation (Anderson et al., 2002; Anderson et al., 1997; Gawlik et al., 2009), but this information is not known for TgSUB2. The activity of subtilisin proteases on their substrate, however, may need neutral or slightly alkaline pH (Shinde et al., 1999). We tested the maturation of TgSUB2 in the *iΔvha1*+ATc mutants and interestingly, we found a significant difference in the ratio between the mature and immature forms of TgSUB2 when compared with the same ratio obtained from the *iΔvha1*-ATc (**Figs. 7C, D**). These results demonstrate that the mature SUB2 is reduced in the *iΔvha1*+ATc mutants, however, the effect on the proteolytic function of TgSUB2 remains to be seen.

Aspartic proteases require a low pH environment for maturation and activity (Darke et al., 1989; Szecsi, 1992). Recently, it was reported that aspartic protease 3 (ASP3) was the maturase for microneme and rhoptry proteins in *T. gondii* (Dogga et al., 2017). With the aim to investigate if maturation of ASP3 was affected in the *iΔvha1-HA* cells, we transfected a plasmid that contained the cDNA of ASP-3Ty, targeted to the *Toxoplasma* UPRT locus into the *iΔvha1* line. Addition of ATc to these cells will ablate the expression of *vha1* and we performed westerns of anti-Ty to follow the maturation of ASP3 (**Fig 7E, F**). There was a significant increase in the level of immature form of the ASP3 in the *iΔvha1-HA+ATc cells* compared to the level present in the *iΔvha1-HA-ATc* (**Fig 7F**). Both cells showed a noticeable level of the immature and mature forms (**Figs. 7E**). It is very likely that the lower levels of mature ASP3 results in a decreased activity because of the more neutral or alkaline conditions of the ELCs. The localization of ASP3 was reported to be the ELCs (Dogga et al., 2017), so we performed IFAs to determine if ASP3 and the V-H^+^-ATPase occupied the same compartment. In dividing intracellular tachyzoites, we observed that ASP3 colocalized with proROP4 and the labeling of the VHa1 encircled the signal of proROP4 (**Fig. 7G**, *dividing PV*). These results indicate that it is very likely that the proton gradient generated by the V-H^+^-ATPase in the immature rhoptries would be responsible for the generation of the ideal environment required for ASP3 maturation into its optimally proteolytically active product (**Fig. 7G**, *dividing PV*). Our results show the role for the V-H^+^-ATPase in regulating the pH of endosomal compartments where important maturation of critical secretory proteins occur.

## DISCUSSION

The function of the V-H^+^-ATPase complex is to pump protons against a concentration gradient and this activity impacts the physiology of every eukaryotic cell (Saroussi and Nelson, 2009). The gradient generated by the activity of the V-H^+^-ATPases in organelles and in membranes of eukaryotic cells is used as a driving force for a number of secondary transport processes (Beyenbach and Wieczorek, 2006). V-H^+^-ATPases are ATP-dependent proton pumps that localize to a variety of cellular membranes in eukaryotic cells (Toei et al., 2010) including endosomes, lysosomes, Golgi-derived vesicles, secretory vesicles and, in some cells, also the plasma membrane (Forgac, 2007). The activity of V-H^+^-ATPases in the endocytic pathway results in a pH gradient that decreases from pH ~6.0 in early endosomes to pH 5.0–5.5 in lysosomes (Hurtado-Lorenzo et al., 2006). Intra-endosomal acidification is required for the enzymatic activity of hydrolytic enzymes (Nishi and Forgac, 2002) among other functions.

The *Toxoplasma* genome shows evidence for the presence of most subunits of the V-H^+^-ATPase with two *a* subunit isoforms, *a1* and *a2* (TgGT1_232830 and TgGT1_290720). In this work we characterized the *a1* subunit, which we termed VHa1. Expression of the *Toxoplasma vha1* gene partially complemented the growth of the mutant strain *Saccharomyces cerevisiae vph1/Δstv1* at pH 7 (Manolson et al., 1994). This was an important result because it showed that VHa1 functions as part of the V-H^+^-ATPase complex and the function of the V-H^+^-ATPase complex could be studied by manipulating the expression of the *vha1* gene. The expression of the *a2* gene in the same strain of *Saccharomyces* also partially complemented their growth indicating that *a2* may also function as part of the complex. We focused on the characterization of *a1* because it showed higher homology to other a subunits of V-H^+^-ATPases of yeast, plants, and mammalian cells and also because the sequence of *a2* contained an extra domain of 12-15 kDa toward the C-terminus, the domain important for proton translocation, which is not found in yeast, plants, or mammalian cells. In addition, our attempts to C-terminally tag VHa2 were not successful.

The V-H^+^-ATPase of *T. gondii* localized to the plasma membrane where it pumps protons outside of the cell, and this activity acted in the recovery of normal cytosolic pH from acid loads. The active proton extrusion into the surrounding media shows that the V-H^+^-ATPase is fully functional at the plasma membrane and protons generated from normal metabolic functions, or artificially induced with acid load, are pumped outside the cell. It is likely that other mechanisms pump protons outside the cell, like a P-type ATPase or a sodium/proton exchanger (Arrizabalaga et al., 2004) thus explaining why parasites are still alive at 48 h even though the absence of VHa1 expression. The localization of the V-H^+^-ATPase to the plasma membrane and its role in regulating intracellular pH was also shown for another apicomplexan, *Plasmodium falciparum* (Hayashi et al., 2000).

The proton gradient generated by the activity of the V-H^+^-ATPase at the plasma membrane contributed to the generation of a membrane potential and mutant parasites were slightly depolarized (Moreno et al., 1998). This depolarization is modest because, most likely, parasites would not survive if their membranes were considerably depolarized. Changes in membrane potential are often linked with ion influx or efflux (Åkerman, 1978).

The expression of the *vha1* gene could be regulated with ATc in the conditional mutants *iΔvha1*. These parasites showed a strong growth defect and strong phenotypic differences, already observed 48 h after starting ATc treatment and even stronger differences were observed when cells were grown longer with ATc. All of the major steps of the lytic cycle were defective, including maturation and secretion of micronemes, invasion, motility, and egress, supporting an essential role for the V-H^+^-ATPase activity in the lytic cycle of *Toxoplasma*. The invasion defect was evident two days after downregulating the expression of VHA1, which was earlier than the microneme secretion defect, which is only evident three days after ATc treatment. The proton pumping activity is already non-functional at day 2 of ATc treatment so it is likely that the invasion decrease at this stage is due to a defect in signaling, which is linked to proton gradients generated by the V-H^+^-ATPase (Roiko et al., 2014).

In addition, intracellular replication was affected in the *Δvha1+ATc* mutants and our interpretation is that intracellular parasites are metabolically active and as such they likely produce large quantities of acid, which needs to be extruded by the active pumping mechanisms at the plasma membrane. We measured intracellular pH in extracellular parasites and that did not appear to be affected when cells were exposed to regular changes in extracellular pH. However, when the parasite cytosol was challenged with propionic acid the recovery response observed in wild type parasites was defective in the *Δvha1+ATc* mutants. This supports an important function for the pump in protecting the cytoplasmic pH and when this activity is missing results in alterations of intracellular pH which will affect cell fitness and replication.

Secretory organelles, like rhoptries and micronemes, are made de novo in daughter parasites during the process of endodyogeny, the mechanism by which *Toxoplasma* replicates (Black, Michael W. and Boothroyd, John C., 2000). A large body of evidence indicates that the biogenesis of rhoptries and micronemes occurs at the Golgi (Tomavo et al., 2013). Trafficking of cargo to the micronemes involves the participation of endosome-like compartments and their proteolytic maturation involves the participation of specific maturases (Parussini et al., 2010) (Dogga et al., 2017). pro-MICs have been observed in structures bearing late endosomal markers (Harper et al., 2006) and nascent micronemes have been observed in close proximity to the plantlike vacuole (PLV) (an endosome-like compartment). Specific inhibitors of the V-H^+^-ATPase proton pump, like bafilomycin A1, reduced the maturation of cathepsin L, an enzyme that localizes to the PLV, that was shown to be self-processed (Dou et al., 2013) and to be involved in the maturation of microneme proteins (Parussini et al., 2010).

Disruption of the activity of the V-H^+^-ATPase negatively impacted the maturation of MIC3 and M2AP. Defective maturation of M2AP leads to miss-targeting of MIC2 as it was shown that proteolytic stabilization of the TgMIC2–M2AP complex within the micronemes is important for the correct packaging within the micronemes and favor its rapid secretion onto the parasite surface (Harper et al., 2006). The localization of M2AP, MIC3, proM2AP and MIC2 were altered in *Δvha1* mutant parasites and microneme organelles were not correctly oriented. **Fig. 8A** proposes a model for the role of the V-H^+^-ATPase in the maturation of micronemes. The model depicts two potential endosome acidic compartments, one expressing higher levels of the V-H^+^-PPase (VP1 compartment as in (Harper et al., 2006)) (Liu et al., 2014) and a second one where the V-H^+^-ATPase predominates (this could be the PLV or VAC). The acidification of these endosomal organelles could be important for either the maturation process of relevant maturases and/or for their specific proteolytic activity on their substrates, as for example essential adhesins secreted for invasion. The presence of more than one of these compartments could represent the means by which lytic activity is regulated by limiting and/or allowing contact with substrates. VHa1 depletion disrupted the biogenesis of the PLV and this resulted in mistargeting of important secretory proteins.

**FIGURE 8:**
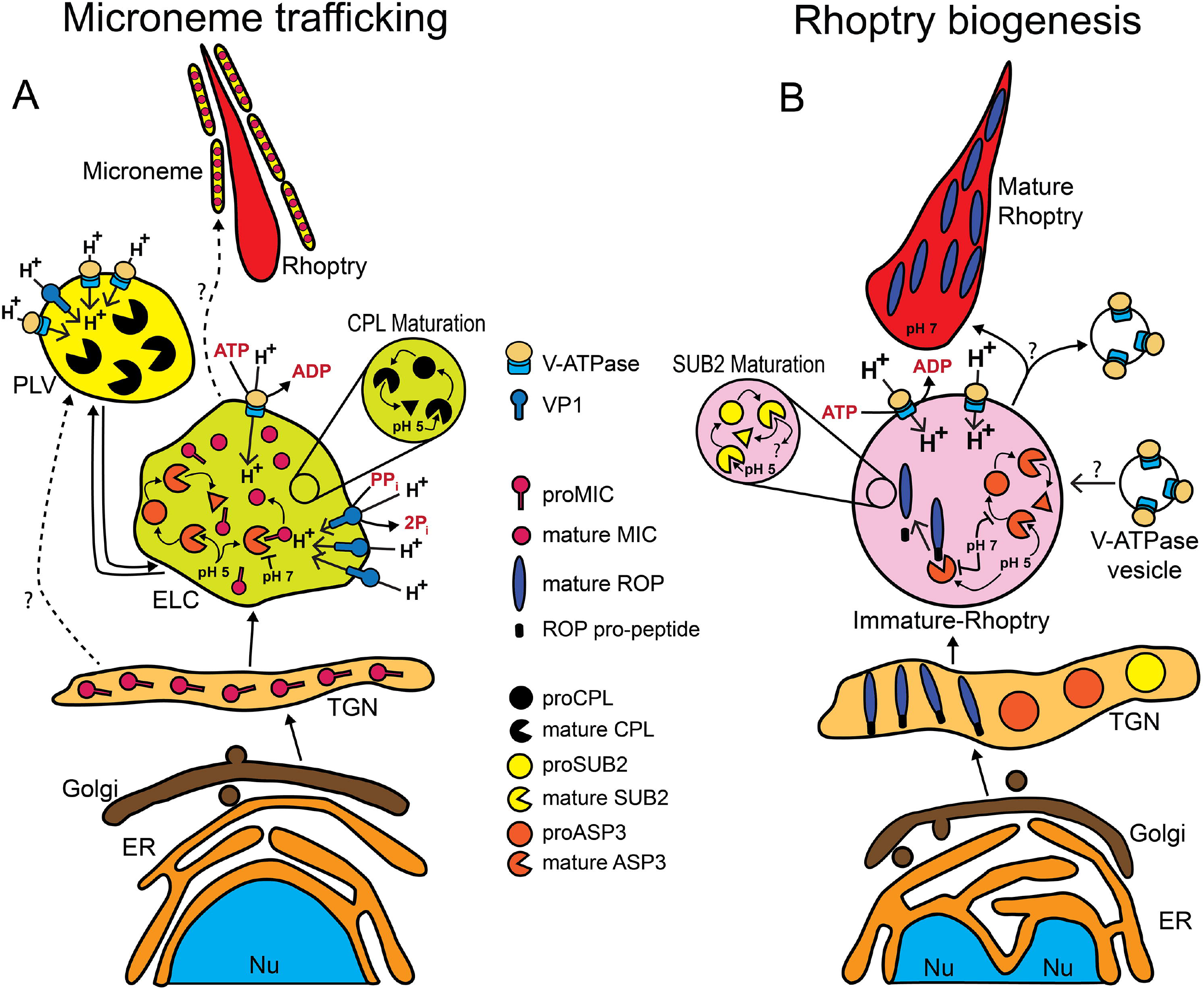
Model of the role of acidification in the maturation of micronemes and rhoptries. A) Microneme biogenesis: Newly synthesized MICs are transported from the Golgi to the TGN and then to the ELC. The ELC is acidified by the vacuolar pyrophosphatase (VP1) and by the V-H^+^-ATPase. The PLV is mainly acidified by the V-H^+^-ATPase and we propose that this compartment would be more acidic and would most likely not contain MICs. Maturation of CPL occurs by self-cleavage and may occur more optimally at the ELC. Maturation of ASP3 occurs in the ELCs. In the V-H^+^-ATPase acidified endosome, proMICs encounter mature ASP3 and/or other lytic enzymes. Cleavage of pro-peptides may occur at this point and once the late endosome is matured, processed MICs would be transported to the microneme organelle. B) Rhoptry biogenesis: immature ROP proteins, immature SUB2, and ASP3 traffic through the TGN. The V-H^+^-ATPase associates with the forming rhoptry and creating the acidic environment that is important for SUB2 and ASP3 maturation. In the immature rhoptries, APS3 would be activate and able to process immature ROPs. According to our model the activity of SUB2 would likely be decreased in low pH. The V-H^+^-ATPase separates by an unknown mechanism allowing the pH of the mature rhoptry to become more neutral, allowing the rhoptry organelle to mature. TGN, *trans*-Golgi-network; PLV, plant-like vacuole; ER, endoplasmic reticulum; Nu, nucleus; ELC, endosome like-compartment.

Rhoptries are club shaped secretory organelles with a discrete neck and bulb region uniquely present in Apicomplexan parasites. Mature rhoptries become club-shaped when mature and they secrete proteins during the process of host cell invasion, an essential function for parasite virulence (Dubremetz, 2007). A number of rhoptry proteins are secreted during host cell invasion and participate in the building of the parasitophorous vacuole in which the parasite will replicate. In addition, some rhoptry proteins are targeted to the host cell nucleus to control host functions (Dubremetz, 2007). In intracellular parasites, the V-H^+^-ATPase labeled immature rhoptries, which have been previously shown to be acidic (pH 3.5-5) in comparison to the more neutral mature rhoptries (pH 5-7) (Shaw et al., 1998). Pro-rhoptries form transiently between the Golgi and the apical area just prior to cytokinesis (Dubremetz, 2007) but the mechanism by which these vesicles form is not known (Ngô et al., 2003). How proteins are delivered to the rhoptries is also not clear, however adaptins have been implicated in this trafficking (Ngô et al., 2003; Venugopal et al., 2017). Vps9, a Rab5 GTP-exchange factor, was shown to be important for ROP protein maturation and processing (Sakura et al., 2016).

Many of the proteins destined for the bulb of the rhoptries (ROPs) contain an N-terminal ER signal peptide and an N-terminal pro-peptide that is cleaved during rhoptry maturation. It was proposed that the N-terminal ROP pro-peptides could be cleaved by subtilisin-like proteases, like TgSUB2, at the SΦX(E/D) motif first identified in ROP1 (Miller et al., 2003; Turetzky et al., 2010). Subtilisin-like proteases typically undergo autocatalytic cleavage of a pro-peptide that initially helps with folding, but must be cleaved before proteolytic activity can occur on substrate proteins (Ikemura and Inouye, 1988; Ikemura et al., 1987). The autocatalytic cleavage and substrate protease activity of subtilisin-like proteases typically have different pH optimums. In the case of subtilisin E from *Bacillus subtilis*, the optimal pH for autocatalytic cleavage of the pro-peptide is pH 7.0, whereas the optimal proteolytic activity on substrate proteins is pH 8.5 (Shinde et al., 1999). TgSUB2 has an atypical cleavage site, which occurs after an acidic residue instead of a basic residue, raising the possibility that the optimal conditions for its cleavage activity are also atypical. There is no experimental evidence to support TgSUB2 maturation in the neutral environment of the ER (Miller et al., 2003).

Aspartic protease 3 (ASP3), a resident aspartyl protease of the ELC, was shown to be critical for invasion and egress of *Toxoplasma* albeit did not appear to be implicated in parasite replication, gliding motility and attachment (Dogga et al., 2017). It was also proposed that the activity of ASP3 superseded the previously proposed roles for TgSUB2 (Miller et al., 2003) and CPL (Parussini et al., 2010) for the maturation of rhoptries and microneme proteins respectively The acidification role of the V-H^+^-ATPase is relevant for the efficient maturation of SUB2 and also ASP3. We propose the model presented in **Fig. 8B**, in which acidification of pro-rhoptries by the V-H^+^-ATPase, would affect the maturation and/or activity of ASP3 resulting in defective maturation of ROP proteins. It is possible that the SUB2 autocatalytic cleavage occurs in the immature rhoptries and low pH, a result of the proton pumping activity of the V-H^+^-ATPase, is required for the cleavage to occur. Acidification of immature rhoptries may also affect maturation of SUB2, but the role of this enzyme in ROP proteins secretion and maturation is less clear. It is evident that the phenotype of the *iΔvha1 (+ATc)* parasites partly mimics the phenotypes of *iΔASP3(+ATc)* on maturation of rhoptries and micronemes. However, *iΔvha1 (+ATc)* parasites also present other phenotypic features probably linked to the role of the pump at the plasma membrane.

This work is the first characterization of the V-H^+^-ATPase protein complex at the molecular level in *Toxoplasma gondii*. The work revealed interesting and novel features of this complex such as its distinct localization in intracellular and extracellular parasites and its functionality at the plasma membrane and the endosomal system. The proton pump function impacts important features of the parasite lytic cycle like egress, replication, as well as maturation of secretory proteins important for invasion of the host cell. The V-H^+^-ATPase proton pump activity supports the processing/synthesis of microneme and rhoptry proteins, which are critical for the lytic cycle of *Toxoplasma*, a central feature of the pathogenesis of the parasite. Specially intriguing is its localization with immature rhoptries and downstream dissociation from rhoptries likely to allow the maturation of the organelle. The traffic of VHa1 to the PLV, soon after egress of tachyzoites and its functional role at the lysosomal-like organelle where it acts in the acidification and maturation of secreted proteins, is also an intriguing feature to adjust to the needs of extracellular tachyzoites. V-ATPases that are no longer needed at the PM are sent to the lysosome for degradation and not to support acidification and maturation of secreted proteins.

In summary, our findings highlight the function of this pump in higher order physiological processes essential for toxoplasma parasitism. In doing so, the activity of the pump impacts the processing/synthesis of virulence factors. Disruption of these delicate mechanisms will alter the most essential aspect of the parasitological cycle of *Toxoplasma*. Our work directly connects the proton pumping activity of the V-H^+^-ATPase complex to processes necessary for *Toxoplasma* virulence.

## MATERIALS AND METHODS

### Chemicals, reagents, and cell cultures

2h, 7h-bis-(2-carboxyethyl)-5(6)-carboxyfluorescein (BCECF) and the acetoxymethyl form (BCECF-AM) were obtained from Molecular Probes. Anhydrotetracycline (ATc), nigericin, 5-fluorodeoxyuridine (FUDR), isopropyl β-D-1-thiogalactopyranoside (IPTG), bafilomycin, and saponin were from Sigma. hTERT or HFF human fibroblasts cells were maintained in Dulbecco’s modified Eagle medium (DMEM) with 10% cosmic calf serum at 37°C with 5% CO_2_. *TATiΔKu80 T. gondii* were used in this study (Sheiner et al., 2011). Culturing of *T. gondii* was performed as described previously (Moreno and Zhong, 1996).

### Genetic manipulation: Epitope-tagging, inducible knockdown, and complementation

Carboxy-terminus tagging was done in the parental line RHTatiΔku80 (*TatiΔku80*) (Sheiner et al., 2011) a parasite line that contains the tetracycline-regulated transactivator system that allows conditional expression of genes (Meissner et al., 2001) and also in which the *ku80* gene was deleted increasing efficiency of homologous recombination (Fox et al., 2009). Primers 1-6 (**Table S1**) were used to create C terminal insert fragments of genes *vha1* (TgGT1_232830), *vhE* (TgGT1_305290), and *vhG* (TgGT1_ 246560) (**Table 1**) that were suitable for cloning into a pLIC-3XHA, pLIC-mNeon-Green, or pLIC-GFP plasmids. Linearized plasmids were transfected into *TATiΔku80* cells followed by drug selection. Upstream gene locus primers 7-9 (**Table S1**) and a pLIC reverse primer (primer 10) were used to verify proper insertion into the correct gene locus. Western blot analyses with rat-α-HA (1:200) confirmed the 3xHA tagging. For tdTomato expressing cell lines, parasites were transfected with a tdTomato overexpressing plasmid (Chtanova et al., 2008) (a gift from Boris Striepen, University of Georgia), enriched using a Bio-Rad S3 cell sorter, and subcloned. For *vha1* knockdowns, a tetracycline-regulatable element was inserted upstream of the translational start codon via double homologous recombination as previously described (Sheiner et al., 2011). Primers 11-14 (**Table S1**) were used to generate the upstream UTR and gene fragments respectively for promoter insertion. Primes 15-18 were used to confirm insertion into the correct genetic locus. Repression of *vha1* was accomplished with 0.5 mg/mL anhydrotetracycline. Complementation was achieved by cloning *vha1* cDNA, using primers 19 and 20 (**Table S1**), into a UPRT cDNA shuttle vector, which contains the 5’ and 3’ UTR’s of the *UPRT* gene (Donald and Roos, 1995). The plasmid was transfected into *iΔvha1-HA* cells expressing tdTomato and selected using 5 μM 5-fluorodeoxyuridine (FUDR) as previously described (Donald and Roos, 1995). Primers 18 and 21 (**Table S1**) were used to verify if *vha1* cDNA inserted into the UPRT gene. C-terminal Ty1 or myc tagging of aspartic protease 3 (ASP3; TgGT1_246550) was performed in the *iΔvha1-HA* strain by co-transfecting a plasmid that contained *CAS9* and a protospacer against the *UPRT gene* locus along with a plasmid that contained the *ASP3-3Ty1 cDNA* or a pLIC plasmid with a C-Terminal myc tag, both plasmids generous gifts from Dominique Soldati-Favre (Dogga et al., 2017). Correct ASP3 C-terminal tags were confirmed by PCR (Primers 26, 27, and/or 10) and western blots with anti-Ty1 or anti-myc.

### Immunofluorescence and western blot analyses, and immuno-electron microscopy

Immunofluorescence assays and immunoblots were performed as described (Liu et al., 2014). IFA images were taken with an Olympus IX-71 inverted fluorescence microscope with a Photometrix CoolSnapHQ CCD camera driven by DeltaVision software or with a Zeiss ELYRA S1 (SR-SIM) Super-resolution microscope. Images were deconvolved using Applied Precision’s Softworx imaging suite using 10 cycles of enhanced ratio deconvolution or with ZEN 2011 software with SIM analysis module for SR-SIM images. For all immunofluorescence analyses (IFA), a control of *TATiΔKu80* with only secondary antibody were used to subtract non-specific background. Immuno-electron microscopy was done using VHa1-HA or VHa1-GFP tagged cells. Parasites were manually lysed and incubated in DMEM media for 0 or 1.5 hours. Cells were then fixed with 4% paraformaldehyde and 0.05% glutaraldehyde for 1 hour on ice and washed once with phosphate buffered saline once. Sample preparation and images were performed by Dr. Wandy Beatty at the Department of Molecular Microbiology, Washington University School of Medicine, St. Louis, MO 63110.

Routine electron microscopy was performed at the Georgia Electron Microcopy Center. Parasites were fixed in 2.5% glutaraldehyde and 2% paraformaldehyde and post-fixed in 2% osmium tetroxide in 0.1M cacodylate buffer (pH 7.2). Samples were then En bloc stained in 0.5% uranyl acetate and dehydrated in an ascending ethanol series and embedded in Spurr’s plastic (Spurr, 1969). Cells were polymerized in an Eppendorf tube at 70 degrees Celsius for 12 hours. Embedded cells were sectioned on grids post-stained with uranyl acetate and lead nitrate. Grids were viewed in a JEOL JEM-1011 transmission electron microscope operated at 80kV.

Quantification was performed following the criteria: micronemes must be at least 2 μm from apical tip to be considered periphery and only one periphery microneme is needed to confirm a positive localization at the periphery; rhoptries must be greater the 0.5 μm and contain a bulb and neck component to be considered mature. EM slices that contained a minimum of 1 visible microneme or rhoptry were enumerated and slices with no visible microneme of rhoptry were not enumerated. Approximately 59-73 parasites were quantified from 3 independent preps.

### Antibody production

Primers 22 and 23 (**Table S1**) were used to clone the first 327 amino acids of the *vha1 gene* into the bacterial inducible expression plasmid pQE-80L. Expression was induced with 1 M Isopropyl β-D-1-thiogalactopyranoside (IPTG) for 2 hours at 37°C. Bacterial lysate was purified with a Thermo scientific HisPur Ni-NTA chromatography cartridge. Mice were injected with 100 μg peptide in Freund’s complete adjuvant and boosted twice every 2 weeks with 50 μg peptide and Freund’s incomplete adjuvant (Chasen et al., 2017). 10-week serum was tested comparing *vha1-HA* lysate with anti-HA and anti-VHa1 to confirm correct size and antibody purity (~37 kDa). The serum was affinity purified and the anti-VHa1 antibody used in IFAs of intracellular and extracellular tachyzoites for localization studies. Work with mice was carried out in strict accordance with the Public Health Service Policy on Humane Care and Use of Laboratory Animals and Association for the Assessment and Accreditation of Laboratory Animal Care guidelines. The animal protocol was approved by the University of Georgia’s Committee on the Use and Care of Animals (protocol A2015 02-025-R2). All efforts were made to humanely euthanize the mice after collecting blood.

### Lytic cycle assays

Plaque assays were performed with confluent hTERT host cells in six-well plates infected with 200 parasites per well and incubated with or without ATc. After 8 days of incubation, parasites were fixed and stained as previously described (Liu et al., 2014). Red/Green invasion assay was performed as described (Kafsack et al., 2004) with some modifications. Freshly egressed parasites were resuspended in ice cold invasion media at a concentration of 2 x 10^7^ ml and incubated on ice for 20 min. The plate was then incubated at 37°C for 5 min, washed with PBS twice, fixed with 3% paraformaldehyde, and blocked with 10% fetal bovine serum (FBS) for 20 min. Rabbit anti-SAG1 polyclonal antibody (1:1000) was used before permeabilization and anti-SAG1 monoclonal antibody (1:500) was used after permeabilization. Counting of red and green labeled parasites was compiled from three independent experiments by counting ten fields of view selected at random. To test for a replication phenotype, tdTomato-expressing parasites were incubated with or without ATc for 24 hours. These parasites (1 x 10^5^) were used to infect (37°C for 30 min) sub-confluent HFF cells previously grown on coverslips. Following this incubation, parasites were washed twice to remove any non-invaded cells and incubated with or without ATc for an additional 24 hours. After 48 hours of total ATc incubation, Toxoplasma-infected cells were washed twice with PBS and fixed with 3% paraformaldehyde for 10 min. Between 115-150 parasitophorous vacuoles (PV) per condition were counted and the number of parasites inside each PV enumerated. These experiments were repeated a minimum of three times. Growth was assayed by infecting confluent hTERT cells in 96 well plates with 4,000 tdTomato-expressing parasites with or without ATc in the media. Parasites were allowed to grow in DMEM media without phenol red for 6 days. A standard curve was used to correlate parasite numbers, (1 x 10^6^ to 1 x 10^1^) with fluorescence values. Egress was stimulated with nigericin as described previously with some modifications (Fruth and Arrizabalaga, 2007). Briefly, tdTomato-expressing parasites were incubated with ATc for 2 days and only PV’s with 2 or more parasites were counted. An additional egress assay was performed using 0.01% saponin. HFF cells seeded onto 35 mm MatTek culture dishes and infected with tdTomato-expressing parasites were pre-incubated for 24 hours with ATc, harvested, and passaged to confluent ibidi culture dishes with ATc. After an additional 24 hours (48 hours total with ATc), parasites were assayed using time-lapse microscopy. Parasites were equilibrated for 2 minutes before adding 20 μL of a 0.01% saponin solution to the dish. A minimum of 6 PV’s per field of view were assayed and the time for each PV to egress was enumerated. If no PV’s egressed after 20 minutes from time of saponin addition, those PVs were assigned an egress time of 1,200 sec. Motility was assayed by resuspending parasites in ringer buffer without calcium with 100 *μ*M EGTA. tdTomato-expressing parasites incubated with or without ATc were tested. 24 hours prior to the experiment, 35 mm MatTek dishes were incubated with 10% FBS to provide sufficient protein to allow a surface conducive for motility. MatTek dishes were washed once with PBS and loaded with 2 mL of Ringer without Ca^2+^. MatTek dishes were chilled on ice and parasites were added and allowed to equilibrate for 1.5 min. Dishes were then placed in the Zeiss LSM 710 Confocal Microscope environmental chamber set to 37°C. 1.8 mM Ca^2+^ was added at the indicated time and motility was quantified as previously described (Fazli et al., 2017). Length of trials was manually traced in ImageJ and the length of parasite movement is reported in μm.

### Yeast complementation

The *a* subunit deficient yeast *(vph1Δstv1Δ)* was used to test complementation (Perzov et al., 2002). Primers 24-25 and 28-29 (**Table S1**) were used to clone *vha1-HA* or *vha2* cDNA, respectively, into the pYES2NT/c yeast galactose inducible plasmid (O’Brien et al., 2015). For inducible expression analysis, yeasts were grown on Complete Supplement Mixture medium lacking uracil (CSM-ura) media at pH 5.5 with 2% galactose or glucose. Harvested yeast lysate was run on a 10% SDS-PAGE gel with rat anti-HA (1:200) used to determine the presence or absence of VHa1 during expressing or non-expressing conditions. To test for complementation, strains were grown on CSM-ura media at pH 7.0 with 2% galactose for 48 (liquid) or 96 hours (plates) at 30°C (O’Brien et al., 2015). For growth in liquid CSM-ura, yeast strains were normalized to an OD_600_ of 1 and diluted 1:10 in liquid media in triplicate per 3 independent trials. Plates were read in a BioTek Synergy H1 hybrid tester every hour for 48 hours under high orbital shaking incubated at 37°C. For growth on plates, yeast strains were normalized to an OD_600_ of 1 and serially diluted 1:10 onto the plates for three independent trials (O’Brien et al., 2015). Plates were photographed 96 hours after incubation. GFP tagging Vph1p and VHa1 was performed using yeast gap repair as previously described (O’Brien et al., 2015).

### pHi, pH_e_, lysotracker, and membrane potential

Internal pH (pH_i_) on *iΔvha1 and iΔvha1–CM* with or without ATc was determined fluorometrically by loading parasites with BCECF-AM as described previously (Moreno et al., 1998). Briefly, cells at a final density of 1 x 10^9^ were loaded with 9 μM BCECF-AM in BAG with 1.5% sucrose and incubated at 37°C for 20 min. Parasites were washed twice with BAG and resuspended to a final density of 1 X 10^9^ cells/ml in BAG and were kept on ice and protected from light. For fluorescence measurements, a 50 μl aliquot of the chilled cell suspension was diluted into 2.45 ml of standard buffer (135 mM NaCl, 5 mM KCl, 1 mM MgSO_4_, 1 mM CaCl_2_, 5 mM glucose and 10 mM Hepes/Tris, pH 7.4) to a final density 2 x 10^7^ cells/ml. The cell suspension was allowed to equilibrate for 2.5 min. in a cuvette before being placed in a Hitachi F-7000 spectrofluorometer. Fluorescence ratios were calculated with excitations at 505 and 440 nm and emission at 530 nm. A standard curve was created using parasites in high potassium standard buffer (140 mM KCl, 1 mM MgSO_4_, 1 mM CaCl_2_, 5 mM glucose and 10 mM Hepes/Tris, pH 7.4) at pH’s ranging from 5.5 to 8 (in 0.5 pH increments) and by adding 5.2 μM of nigericin to the cell suspension. To determine the effect of changing extracellular pH (pH_e_) on pH_i_, *iΔvha1* with or without ATc were loaded with BCECF-AM as described previously (Moreno et al., 1998) and incubated in standard buffer at varying pH_e_. Proton extrusion was determined with BCECF free acid as described previously (Pace et al., 2011). Briefly, a 100 μl aliquot of a cell suspension (at 1 x 10^9^ cell/mL) was diluted into 2.45 ml in a weakly buffered solution (0.1 mM HEPES-Tris) with 0.38 μM BCECF free acid. A standard curve of known pH’s was used to determine the change of the extracellular pH. The *VHa1* gene was C-terminally tagged with mNeon-green using the same genetic strategy used for HA-tagging. Clonal populations were isolated and were loaded with 10 μM LysoTracker red for 30 min at 37°C in BAG, washed twice with BAG, and placed in 35 mm MatTek dishes in Ringer buffer (155 mM NaCl, 3 mM KCl, 1 mM MgCl_2_, 3 mM NaH_2_PO_4_, 2 mM CaCl_2_, and 10 mM Hepes, and 10 mM dextrose). For quantification of Lysotracker stained compartments, the number of vacuoles from 10 fields of view with at least 5 parasites was tallied. To minimize bias, counting was performed double blinded. Membrane potential was measured fluorometrically with bisoxonol (Moreno et al., 1998). Bisoxonol (0.2 μM) was added to 5 x 10^7^ cells suspended in 2.5 ml of standard buffer at pH 7.5 or 7.0 at 37°C with excitation at 540 nm and emission at 580 nm. The last 100 seconds of the tracing were averaged at each pH and the ratio of pH 7.5/7.0 determined.

### Microneme secretion and maturation

Microneme secretion was determined as described previously (Liu et al., 2014). M2AP maturation was measured by using an antibody created against the propeptide part (a generous gift from Vern Carruthers, University of Michigan) (Harper et al., 2006). *iΔvha1-HA* parasites were incubated with ATc for 0 to 3 days, lysed, and run on an 10% SDS-PAGE gel and probed with anti-proM2AP at 1:500 and anti-Tubulin. Immunoblots were probed with either antiM2AP or anti-MIC and analyzed using pixel density by ImageJ or Image Studio (where indicated). Quantification was determined either by a ratio of immature/mature or mature/tubulin. Maturation of CPL was performed using an antibody against CPL, which recognizes both the mature and immature forms. *iΔvha1-HA* parasites were incubated with ATc for 0 to 3 days, lysed, and run on an 10% SDS-PAGE gel. Anti-CPL at 1:500 was used to identify proCPL from mature CPL. Pixel densities generated using ImageJ of proCPL and mature CPL were normalized to tubulin and the tubulin normalized ratios were used to determine the ratio proCPL/CPL.

### Maturation of Rhoptry Proteins and Quantification of Immature Rhoptries

Maturation of rhoptry proteins was analyzed using α-ROP4 (UVT-68, Rabbit 1:500), α-ROP7 (1:1000), and α-TgCA_RP (Guinea Pig 1:1000). Parasites were harvested and analyzed as described above after 0 and 3 days of ATc treatment. Quantification of band intensity was performed using signal intensity determined by LI-COR Image Studio Software. Percentage of immature protein was determined by dividing the band intensity of the immature band by the total combined intensity of mature and immature bands. Immunofluorescence assays of *iΔvha1-HA* tachyzoites grown in without ATc or 3 days with Tc were performed using the immature rhoptry specific antibody α- proROP4 (UVT-70, 1:500), rhoptry bulb antibody α-TgCA_RP (Guinea Pig 1:1000). α-IMC1 (Mouse, 1:500) was used to identify dividing tachyzoites. For each experiment 100 vacuoles (containing 2-8 tachyzoites) were examined for the presence or absence of tachyzoites containing nascent rhoptries (α-proROP4 labeling) and/or undergoing cytokinesis (α-IMC1 labeling of daughter cells).

### Statistical analysis

All statistical analyses were performed using GraphPad Prism 6. Unless otherwise noted, all error bars are presented as the standard error of the mean (SEM) and from a minimum of three independent trials. Significant differences were only considered if *P* values were < 0.05.

## ACKNOWLEDGMENTS

We thank Vern Carruthers for the anti-MIC2, proM2AP and CPL antibodies, Maryse Lebrun for the anti-MIC3 antibody, and Kami Kim for the anti-TgSUB2 antibody. We thank Dominique Soldati-Favre for sending us plasmids for C-terminal tagging of ASP3. We thank Melissa Storey for their work in creating the VHa1 antibody. We also thank the Georgia Electron Microcopy Center for assisting with the high-quality electron microscopy images and the Biomedical Microscopy Core of the Coverdell Center for the use of their microscopes. Wandy Betty from Washington University, St Louis performed the immunoelectron microscopy presented in Figure 1. We thank Zhicheng Dou, Drew Etheridge and Vern Carruthers for insightful suggestions and for reading the manuscript. Funding for this work was provided by the U.S. National Institutes of Health (grants AI-096836 to SNJM and AI-100913 to VJS). NC was supported by a pre-doctoral fellowship from the American Heart Association (14PRE19100003). AJS was partly funded through a UGA OVPR fellowship as part of the NIH’s T32 training grant T32AI060546.

## AUTHOR CONTRIBUTIONS

A.J.S. performed and designed most of the experiments, analyzed the data, and wrote the manuscript. N.M.C. performed the rhoptry phenotype characterization. E.J.D. assisted with the pH and proton extrusion measurements and helped with the characterization of the mutant phenotype. S.A.V. performed the super resolution images and helped with the motility assay. B.A. performed and designed the routine electron microscopy experiments and their interpretation. V.J.S. supervised and designed the yeast complementation experiments. S.N.J.M. is the principal investigator who supervised the whole project.

---

**Supplemental Video 1: pH indicator mCherry-SEpHluorin expressed in the PLV.** *iΔvha1-HA* parasites were transfected with a plasmid that targeted mCherry-SEpHluorin (Koivusalo et al., 2010) to the PLV. Extracellular parasites were placed in MatTek dishes with Ringer buffer at pH 7.3. At 15 seconds, 20 mM NH_4_Cl was added. Due to the acidic nature of the PLV, the green channel (which responds to changes in pH) was not seen until after alkalization, confirming that the PLV is normally acidic. The change in fluorescence indicates that the indicator is functional.

**Supplemental Table 1:**

Primer sequences used in this work

## Supplemental Figure legends

**FIGURE S1: The Vacuolar H^+^-ATPase of *Toxoplasma gondii*.** A) Scheme of the various subunits and domains of the Vacuolar-H^+^-ATPase in *T. gondii*. Structural design adapted from (Forgac, 2007). B) VHa1 sequence modeling using Protter version 1.0, which predicts 7 transmembrane domains. The 27 amino acids that form the predicted signal peptide are marked in red. Scale bar is 5 amino acids. C) Scheme of the primers used to determine correct tag insertion. D) PCR using an upstream gene specific forward primer (Primers 7-9) and reserve pLIC primer (Primer 10). Primer sets specific for *vha1, vhG, vhE* being HA tagged were also compared against a *TATiΔKu80* strain (WT).

**FIGURE S2: The *T. gondii* V-H^+^-ATPase localizes to the plasma membrane and the PLV.** A) Immunofluorescence assays of extracellular tachyzoites treated for 4 hours with alpha toxin as detected using anti-HA anti-IMC1. B) IFAs of *iΔvha1-HA* intracellular parasites with anti-HA and anti-SAG1 in hTERT host cells. C) Super resolution IFA’s using anti-HA and plant like vacuole markers anti-VP1. Arrow points to an internalized puncta that co-localizes with PLV marker VP1. Dashed line indicates the outline of the parasite. D) VHa1-HA parasites expressing *VHG* C-terminally Ty1 tagged. E) IFAs of extracellular parasites expressing VHG-HA with anti-HA and anti-VP1. F) IFAs of extracellular parasites expressing VHE-HA with anti-HA and anti-VP1. G) Quantification of IFA images showing the formation of a large central VHa1 puncta and its co-localization with PLV marker VP1 in function of time. H) Western blots of *TATiAku80* or VHa1-HA lysates probed with anti-VHa1 or anti-HA, respectively. I) IFA of extracellular parasites with anti-VHa1. IFA scale bars are 2 μm.

**FIGURE S3. Yeast complementation and localization of VHa1 in *Δvph1Δstv1* yeast.** A) Spotted serial dilutions of yeast suspensions normalized to an OD of 1 of yeast expressing VHa1 *(left panel)* or VHa2 *(right panel)*. WT, wild type yeast; EV, empty pYES2 vector; *vha1/vha2*, pYES2 vector with *vha1* or *vha2* cDNA. Suspensions were grown on CSM-ura plates with 2% galactose at pH 7.0 for 96 hours. Blots are representative from 3 independent trials done in duplicate. B) cDNA from *vha1* or *vph1* were fused to GFP and transformed into *Δvph1Δstvl* yeast. Arrows point to the yeast vacuole.

**FIGURE S4: Conditional knock down of the *vha1 gene*.** A) Cartoon showing the promoter insertion strategy and localization of primers used for genetic validation. CAT, chloramphenicol acetyltransferase; HA, 3 hemagglutinin epitope tags; T7/S4, tetracycline inducible promoter; DHFR, pyrimethamine resistance cassette; P, endogenous promoter; PS1, primer set 1. B) PCR with PS1 to confirm integration of cassette upstream of the translational start codon. C) IFA’s showing the regulation of expression of *vha1* by ATc. D) Cartoon showing the strategy used for complementation of the *iΔvha1-HA* parasites with *vha1* cDNA into the *uprt* gene locus. Scale bars are 2 μm.

**FIGURE S5. VHa1 plays roles in the lytic cycle of the parasite.** A) Growth of *iΔvha1-HA* parasites which were preincubated for the indicated days, washed twice, and added to fresh host cells without ATc and their growth monitored. Around 100 tDtomato expressing parasites were put into confluent hTERT cells in a 96 well plate and allowed to grow for 8 days. Data are represented as the average of three independent trials done in triplicate. B) Trypan exclusion of *iΔvha1-HA* parasites incubated with or without ATc for 0 or 3 days. Parasites were either heat treated at 70°C for 5 min or not heat treated and exposed to 1.5% trypan, washed twice, and the number excluding trypan blue enumerated. Data are from three independent trials. C) Growth rate of *vha1-HA, iΔvha1-HA*, and *iΔvha1-HA-CM* tDtomato expressing parasites. Slopes were calculated from days 2-5. Data are from three independent trials. D) *iΔvha1-HA* and *iΔvha1-HA-CM* parasites incubated for 0 (-) or 2 (+) days with ATc were exposed to 0.01% saponin after 120 seconds. 3 independent trails were performed in duplicate where the time to egress after saponin addition was recorded. \**P* < 0.05; \*\**P* < 0.01; \*\*\**P* < 0.001.

**FIGURE S6. Microneme secretion of AMA1.** A) Western blots of total lysates and constitutive secretion (con.) of *iΔvha1-HA* or *iΔvha1-HA-CM* parasites grown with and without ATc. Parasites were incubated for 30 min in invasion media (DMEM-HG with 20 mM HEPES) and supernatant (Con.) and pellet (Lysate) were separated by centrifugation. Immunoblot shows labeling of immature AMA1 (arrow head) and mature AMA1 (arrow) of *iΔvha1-HA* or *Δvha1-HA-CM* parasites grown with and without ATc. GRA1 control is shown in the bottom panel. B) Ratio of AMA1/GRA1 as determined by pixel density using Image Studio from 3 independent trials. Graph error bars are SEM. Con. Means constitutive secretion. Statistical analysis were performed using a one way ANOVA where \**P* < 0.05; \*\**P* < 0.01; \*\*\*\**P* < 0.0001.

**FIGURE S7. Parental controls for microneme and rhoptry maturation.** A) Western blots of total lysates of *vha1-HA* parasites grown with and without ATc showing the staining of proM2AP (arrow head) compared to mature M2AP (arrow) (top) MIC3 (mid panel) and Tubulin control (bottom). B) Ratio of proM2AP/M2AP pixel density. C) Ratio of density of the bands of mature MIC3/Tubulin. D) Western blots of total lysates of *vha1-HA* parasites grown with and without ATc showing normal maturation of ROP4, ROP7, and TgCA_RP. Arrowhead indicates immature protein and arrow denotes mature protein. E) Quantification of bands shown in F by ratio of immature/mature. 3 independent biological replicates were performed. F) Western blots of total lysates of *vha1-HA* parasites grown with and without ATc showing the maturation of CPL. G) Quantification of immature/mature CPL. B-C, E, and G were quantified by pixel density using Image Studio from 3 independent trials. Graph error bars are SEM. Statistical analysis were performed using a Student’s *t* test where NS is not significant.

**FIGURE S8. The localization of MIC2 is altered in *iΔvha1-HA* parasites.** IFAs of *iΔvha1-HA* parasites grown with or without ATc (3 days) using rat-anti-HA (1:25) and anti-MIC2 at a concentration of 1:400. Arrows point to non-peripheral atypical labeling. Scale bars are 2 μm.

**FIGURE S9. The V-H^+^-ATPase associates with pro-rhoptries.** A) Super resolution IFA of *iΔvha1-HA* without ATc fixed 24 hours after invasion with 4% paraformaldehyde. Antibodies against HA, labeling VHa1, (rat-anti-HA, 1:25, green), anti-proROP4 (Mab UVT-70, 1:500, red), and anti-TgCA_RP (Guinea Pig, 1:1000, blue) were used for intracellular localizations. B) Super resolution IFA of intracellular *iΔvha1-HA* clones without ATc using antibodies against HA (1:25), proROP4 (1:500), and TgCA_RP (1:1000). Cells were fixed 24 hours after invasion with 4% paraformaldehyde. C) Super resolution IFA of VHE-HA parasite clones using anti-HA (labeling subunit E) and anti-proROP4 (1:500, red). Cells were fixed 24 hours after invasion with 4% paraformaldehyde. D) Same as C but with VHG-HA clones. Scale bars are 2 μm.

## REFERENCES

Åkerman, K.E.O. (1978). Changes in membrane potential during calcium ion influx and efflux across the mitochondrial membrane. Biochimica et Biophysica Acta (BBA) - Bioenergetics. 502(2), 359–366. DOI: 10.1016/0005-2728(78)90056-7.

Anderson, E.D., Molloy, S.S., Jean, F., Fei, H., Shimamura, S., and Thomas, G. (2002). The ordered and compartment-specific autoproteolytic removal of the furin intramolecular chaperone is required for enzyme activation. Journal of Biological Chemistry. 277(15), 12879–12890. DOI: 10.1074/jbc.M108740200.

Anderson, E.D., VanSlyke, J.K., Thulin, C.D., Jean, F., and Thomas, G. (1997). Activation of the furin endoprotease is a multiple‐ step process: requirements for acidification and internal propeptide cleavage. The EMBO journal. 16(7), 1508–1518.

Arrizabalaga, G., Ruiz, F., Moreno, S., and Boothroyd, J.C. (2004). Ionophore-resistant mutant of *Toxoplasma gondii* reveals involvement of a sodium/hydrogen exchanger in calcium regulation. The Journal of Cell Biology. 165(5), 653–662. DOI: 10.1083/jcb.200309097.

Beyenbach, K.W., and Wieczorek, H. (2006). The V-type H^+^ ATPase: molecular structure and function, physiological roles and regulation. Journal of Experimental Biology. 209(4), 577–589. DOI: 10.1242/jeb.02014.

Black, M.W., and Boothroyd, J.C. (2000). Lytic cycle of *Toxoplasma gondii*. Microbiol Mol Biol Rev. 64(3), 607–623.

Black, M.W., and Boothroyd, J.C. (2000). Lytic Cycle of *Toxoplasma gondii*. Microbiology and Molecular Biology Reviews. 64(3), 607–623. DOI: 10.1128/mmbr.64.3.607-623.2000.

Borges-Pereira, L., Budu, A., McKnight, C.A., Moore, C.A., Vella, S.A., Triana, M.A.H., Liu, J., Garcia, C.R., Pace, D.A., and Moreno, S.N. (2015). Calcium Signaling throughout the *Toxoplasma gondii* Lytic Cycle a study using genetically encoded calcium indicators. Journal of Biological Chemistry. 290(45), 26914–26926.

Carruthers, V.B., Giddings, O.K., and Sibley, L.D. (1999). Secretion of micronemal proteins is associated with toxoplasma invasion of host cells. Cell Microbiol. 1(3), 225–235.

Cérède, O., Dubremetz, J.F., Bout, D., and Lebrun, M. (2002). The *Toxoplasma gondii* protein MIC3 requires propeptide cleavage and dimerization to function as adhesin. The EMBO Journal. 21(11), 2526–2536. DOI: 10.1093/emboj/21.11.2526.

Chasen, N.M., Asady, B., Lemgruber, L., Vommaro, R.C., Kissinger, J.C., Coppens, I., and Moreno, S.N.J. (2017). A Glycosylphosphatidylinositol-Anchored Carbonic Anhydrase-Related Protein of *Toxoplasma gondii* Is Important for Rhoptry Biogenesis and Virulence. mSphere. 2(3), e00027–00017. DOI: 10.1128/mSphere.00027-17.

Chtanova, T., Schaeffer, M., Han, S.-J., van Dooren, G.G., Nollmann, M., Herzmark, P., Chan, S.W., Satija, H., Camfield, K., Aaron, H., et al. (2008). Dynamics of Neutrophil Migration in Lymph Nodes during Infection. Immunity. 29(3), 487–496. DOI: 10.1016/j.immuni.2008.07.012.

Darke, P.L., Leu, C.T., Davis, L.J., Heimbach, J.C., Diehl, R.E., Hill, W.S., Dixon, R.A., and Sigal, I.S. (1989). Human immunodeficiency virus protease. Bacterial expression and characterization of the purified aspartic protease. Journal of Biological Chemistry. 264(4), 2307–2312.

Demaurex, N. (2002). pH Homeostasis of Cellular Organelles. Physiology. 17(1), 1–5.

Dogga, S.K., Mukherjee, B., Jacot, D., Kockmann, T., Molino, L., Hammoudi, P.-M., Hartkoorn, R.C., Hehl, A.B., and Soldati-Favre, D. (2017). A druggable secretory protein maturase of Toxoplasma essential for invasion and egress. eLife. 6, e27480. DOI: 10.7554/eLife.27480.

Donald, R.G., and Roos, D.S. (1995). Insertional mutagenesis and marker rescue in a protozoan parasite: cloning of the uracil phosphoribosyltransferase locus from *Toxoplasma gondii*. Proceedings of the National Academy of Sciences. 92(12), 5749–5753.

Dou, Z., Coppens, I., and Carruthers, V.B. (2013). Non-canonical maturation of two papain-family proteases in *Toxoplasma gondii*. J Biol Chem. 288(5), 3523–3534. DOI: 10.1074/jbc.M112.443697.

Dubremetz, J.F. (2007). Rhoptries are major players in *Toxoplasma gondii* invasion and host cell interaction. Cell Microbiol. 9(4), 841–848. Published online 2007/03/10 DOI: 10.1111/j.1462-5822.2007.00909.x.

El Hajj, H., Papoin, J., Cerede, O., Garcia-Reguet, N., Soete, M., Dubremetz, J.F., and Lebrun, M. (2008). Molecular signals in the trafficking of *Toxoplasma gondii* protein MIC3 to the micronemes. Eukaryot Cell. 7(6), 1019–1028. DOI: 10.1128/EC.00413-07.

Fazli, M.S., Vella, S.A., Moreno, S.N., and Quinn, S. (2017). Computational motility tracking of calcium dynamics in *Toxoplasma gondii*. arXiv preprint arXiv:1708.01871.

Forgac, M. (1989). Structure and function of vacuolar class of ATP-driven proton pumps. Physiological Reviews. 69(3), 765–796.

Forgac, M. (1999). Structure and Properties of the Vacuolar H^+^-ATPases. Journal of Biological Chemistry. 274(19), 12951–12954. DOI: 10.1074/jbc.274.19.12951.

Forgac, M. (2007). Vacuolar ATPases: rotary proton pumps in physiology and pathophysiology. Nat Rev Mol Cell Biol. 8(11), 917–929.

Fox, B.A., Ristuccia, J.G., Gigley, J.P., and Bzik, D.J. (2009). Efficient gene replacements in *Toxoplasma gondii* strains deficient for nonhomologous end joining. Eukaryot Cell. 8(4), 520–529. DOI: 10.1128/EC.00357-08.

Fruth, I.A., and Arrizabalaga, G. (2007). *Toxoplasma gondii*: Induction of egress by the potassium ionophore nigericin. International Journal for Parasitology. 37(14), 1559–1567. DOI: 10.1016/j.ijpara.2007.05.010.

Garcia-Réguet, N., Lebrun, M., Fourmaux, M.-N., Mercereau-Puijalon, O., Mann, T., Beckers, C.J.M., Samyn, B., Van Beeumen, J., Bout, D., and Dubremetz, J.-F. (2000). The microneme protein MIC3 of *Toxoplasma gondii* is a secretory adhesin that binds to both the surface of the host cells and the surface of the parasite. Cellular Microbiology. 2(4), 353–364. DOI: 10.1046/j.1462-5822.2000.00064.x.

Gawlik, K., Shiryaev, S.A., Zhu, W., Motamedchaboki, K., Desjardins, R., Day, R., Remacle, A.G., Stec, B., and Strongin, A.Y. (2009). Autocatalytic activation of the furin zymogen requires removal of the emerging enzyme’s N-terminus from the active site. PLoS One. 4(4), e5031.

Hajagos, B.E., Turetzky, J.M., Peng, E.D., Cheng, S.J., Ryan, C.M., Souda, P., Whitelegge, J.P., Lebrun, M., Dubremetz, J.-F., and Bradley, P.J. (2012). Molecular Dissection of Novel Trafficking and Processing of the *Toxoplasma gondii* Rhoptry Metalloprotease Toxolysin-1. Traffic. 13(2), 292–304. DOI: 10.1111/j.1600-0854.2011.01308.x.

Harper, J.M., Huynh, M.H., Coppens, I., Parussini, F., Moreno, S., and Carruthers, V.B. (2006). A cleavable propeptide influences Toxoplasma infection by facilitating the trafficking and secretion of the TgMIC2-M2AP invasion complex. Mol Biol Cell. 17(10), 4551–4563. DOI: 10.1091/mbc.E06-01-0064.

Hayashi, M., Yamada, H., Mitamura, T., Horii, T., Yamamoto, A., and Moriyama, Y. (2000). Vacuolar H^+^-ATPase Localized in Plasma Membranes of Malaria Parasite Cells, *Plasmodium falciparum*, Is Involved in Regional Acidification of Parasitized Erythrocytes. Journal of Biological Chemistry. 275(44), 34353–34358. DOI: 10.1074/jbc.M003323200.

Hehl, A.B., Lekutis, C., Grigg, M.E., Bradley, P.J., Dubremetz, J.-F., Ortega-Barria, E., and Boothroyd, J.C. (2000). *Toxoplasma gondii* Homologue of *Plasmodium* Apical Membrane Antigen 1 Is Involved in Invasion of Host Cells. Infection and Immunity. 68(12), 7078.

Hurtado-Lorenzo, A., Skinner, M., Annan, J.E., Futai, M., Sun-Wada, G.-H., Bourgoin, S., Casanova, J., Wildeman, A., Bechoua, S., Ausiello, D.A., et al. (2006). V-ATPase interacts with ARNO and Arf6 in early endosomes and regulates the protein degradative pathway. Nat Cell Biol. 8(2), 124–136. DOI: 10.1038/ncb1348

Huynh, M.H., and Carruthers, V.B. (2009). Tagging of endogenous genes in a *Toxoplasma gondii* strain lacking Ku80. Eukaryot Cell. 8(4), 530–539. DOI: 10.1128/EC.00358-08.

Ikemura, H., and Inouye, M. (1988). In vitro processing of pro-subtilisin produced in *Escherichia coli*. Journal of Biological Chemistry. 263(26), 12959–12963.

Ikemura, H., Takagi, H., and Inouye, M. (1987). Requirement of pro-sequence for the production of active subtilisin E in *Escherichia coli*. Journal of Biological Chemistry. 262(16), 7859–7864.

Kafsack, B.F., Beckers, C., and Carruthers, V.B. (2004). Synchronous invasion of host cells by *Toxoplasma gondii*. Mol Biochem Parasitol. 136(2), 309–311.

Koivusalo, M., Welch, C., Hayashi, H., Scott, C.C., Kim, M., Alexander, T., Touret, N., Hahn, K.M., and Grinstein, S. (2010). Amiloride inhibits macropinocytosis by lowering submembranous pH and preventing Rac1 and Cdc42 signaling. J Cell Biol. 188(4), 547–563. Published online 2010/02/17 DOI: 10.1083/jcb.200908086.

Leng, X.H., Nishi, T., and Forgac, M. (1999). Transmembrane topography of the 100-kDa a subunit (Vph1p) of the yeast vacuolar proton-translocating ATPase. J Biol Chem. 274(21), 14655–14661.

Liu, J., Pace, D., Dou, Z., King, T.P., Guidot, D., Li, Z.-H., Carruthers, V.B., and Moreno, S.N.J. (2014). A vacuolar-H^+^-pyrophosphatase (TgVP1) is required for microneme secretion, host cell invasion, and extracellular survival of *Toxoplasma gondii*. Molecular Microbiology. 93(4), 698–712. DOI: 10.1111/mmi.12685.

Manolson, M.F., Wu, B., Proteau, D., Taillon, B.E., Roberts, B.T., Hoyt, M.A., and Jones, E.W. (1994). *STV1* gene encodes functional homologue of 95-kDa yeast vacuolar H^+^-ATPase subunit Vph1p. Journal of Biological Chemistry. 269(19), 14064–14074.

Meissner, M., Brecht, S., Bujard, H., and Soldati, D. (2001). Modulation of myosin A expression by a newly established tetracycline repressor-based inducible system in *Toxoplasma gondii*. Nucleic Acids Res. 29(22), E115.

Miller, S.A., Thathy, V., Ajioka, J.W., Blackman, M.J., and Kim, K. (2003). TgSUB2 is a Toxoplasma gondii rhoptry organelle processing proteinase. Molecular microbiology. 49(4), 883–894.

Miranda, K., Pace, D.A., Cintron, R., Rodrigues, J.C.F., Fang, J., Smith, A., Rohloff, P., Coelho, E., De Haas, F., De Souza, W., et al. (2010). Characterization of a novel organelle in *Toxoplasma gondii* with similar composition and function to the plant vacuole. Molecular Microbiology. 76(6), 1358–1375. DOI: 10.1111/j.1365-2958.2010.07165.x.

Moreno, N.J.S., Zhong, L., Lu, H.-G., Souza, W.D.E., and Benchimol, M. (1998). Vacuolar-type H^+^-ATPase regulates cytoplasmic pH in *Toxoplasma gondii* tachyzoites. Biochemical Journal. 330(2), 853–860. DOI: 10.1042/bj3300853.

Moreno, S.N.J., and Zhong, L. (1996). Acidocalcisomes in *Toxoplasma gondii* tachyzoites. Biochemical Journal. 313(2), 655.

Ngo, H.M., Yang, M., and Joiner, K.A. (2004). Are rhoptries in Apicomplexan parasites secretory granules or secretory lysosomal granules? Mol Microbiol. 52(6), 1531–1541. Published online 2004/06/10 DOI: 10.1111/j.1365-2958.2004.04056.x.

Ngô, H.M., Yang, M., Paprotka, K., Pypaert, M., Hoppe, H., and Joiner, K.A. (2003). AP-1 in *Toxoplasma gondii* Mediates Biogenesis of the Rhoptry Secretory Organelle from a Post-Golgi Compartment. Journal of Biological Chemistry. 278(7), 5343–5352. DOI: 10.1074/jbc.M208291200.

Nishi, T., and Forgac, M. (2002). The vacuolar (H^+^)-ATPases--nature’s most versatile proton pumps. Nat Rev Mol Cell Biol. 3(2), 94–103. DOI: 10.1038/nrm729.

O’Brien, K.M., Lindsay, E.L., and Starai, V.J. (2015). The Legionella pneumophila Effector Protein, LegC7, Alters Yeast Endosomal Trafficking. PLOS ONE. 10(2), e0116824. DOI: 10.1371/journal.pone.0116824.

Pace, D.A., Fang, J., Cintrón, R., Docampo, M.D., and Moreno, S.N.J. (2011). Overexpression of a Cytosolic Pyrophosphatase (TgPPase) Reveals a Regulatory Role of Pyrophosphate in Glycolysis for *Toxoplasma gondii*. The Biochemical journal. 440(2), 229–240. DOI: 10.1042/BJ20110641.

Pace, D.A., McKnight, C.A., Liu, J., Jimenez, V., and Moreno, S.N.J. (2014). Calcium Entry in *Toxoplasma gondii* and Its Enhancing Effect of Invasion-linked Traits. Journal of Biological Chemistry. 289(28), 19637–19647. DOI: 10.1074/jbc.M114.565390.

Paredes-Santos, T.C., de Souza, W., and Attias, M. (2012). Dynamics and 3D organization of secretory organelles of *Toxoplasma gondii*. Journal of Structural Biology. 177(2), 420–430. DOI: 10.1016/j.jsb.2011.11.028.

Parussini, F., Coppens, I., Shah, P.P., Diamond, S.L., and Carruthers, V.B. (2010). Cathepsin L occupies a vacuolar compartment and is a protein maturase within the endo/exocytic system of *Toxoplasma gondii*. Molecular Microbiology. 76(6), 1340–1357. DOI: 10.1111/j.1365-2958.2010.07181.x.

Perzov, N., Padler-Karavani, V., Nelson, H., and Nelson, N. (2002). Characterization of yeast VATPase mutants lacking Vph1p or Stv1p and the effect on endocytosis. Journal of Experimental Biology. 205(9), 1209–1219.

Roiko, M.S., Svezhova, N., and Carruthers, V.B. (2014). Acidification Activates *Toxoplasma gondii* Motility and Egress by Enhancing Protein Secretion and Cytolytic Activity. PLOS Pathogens. 10(11), e1004488. DOI: 10.1371/journal.ppat.1004488.

Sakura, T., Sindikubwabo, F., Oesterlin, L.K., Bousquet, H., Slomianny, C., Hakimi, M.-A., Langsley, G., and Tomavo, S. (2016). A Critical Role for *Toxoplasma gondii* Vacuolar Protein Sorting VPS9 in Secretory Organelle Biogenesis and Host Infection. Scientific Reports. 6, 38842. DOI: 10.1038/srep38842.

Saroussi, S., and Nelson, N. (2009). Vacuolar H^+^-ATPase-an enzyme for all seasons. Pflugers Arch. 457(3), 581–587. DOI: 10.1007/s00424-008-0458-9.

Shaw, M.K., Roos, D.S., and Tilney, L.G. (1998). Acidic compartments and rhoptry formation in *Toxoplasma gondii*. Parasitology. 117(5), 435–443. Published online 1998/11/01 DOI: undefined.

Sheiner, L., Demerly, J.L., Poulsen, N., Beatty, W.L., Lucas, O., Behnke, M.S., White, M.W., and Striepen, B. (2011). A Systematic Screen to Discover and Analyze Apicoplast Proteins Identifies a Conserved and Essential Protein Import Factor. PLOS Pathogens. 7(12), e1002392. DOI: 10.1371/journal.ppat.1002392.

Shinde, U., Fu, X., and Inouye, M. (1999). A pathway for conformational diversity in proteins mediated by intramolecular chaperones. Journal of Biological Chemistry. 274(22), 15615–15621.

Sidik, S.M., Huet, D., Ganesan, S.M., Huynh, M.-H., Wang, T., Nasamu, A.S., Thiru, P., Saeij, J.P.J., Carruthers, V.B., Niles, J.C., et al. (2016). A Genome-wide CRISPR Screen in Toxoplasma Identifies Essential Apicomplexan Genes. Cell. 166(6), 1423–1435.e1412. DOI: 10.1016/j.cell.2016.08.019.

Smardon, A.M., Tarsio, M., and Kane, P.M. (2002). The RAVE Complex Is Essential for Stable Assembly of the Yeast V-ATPase. Journal of Biological Chemistry. 277(16), 13831–13839. DOI: 10.1074/jbc.M200682200.

Soldati, D., Dubremetz, J.F., and Lebrun, M. (2001). Microneme proteins: structural and functional requirements to promote adhesion and invasion by the apicomplexan parasite *Toxoplasma gondii*. International Journal for Parasitology. 31(12), 1293–1302. DOI: 10.1016/S0020-7519(01)00257-0.

Soldati, D., Lassen, A., Dubremetz, J.-F., and Boothroyd, J.C. (1998). Processing of Toxoplasma ROP1 protein in nascent rhoptries. Molecular and Biochemical Parasitology. 96(1), 37–48. DOI: 10.1016/S0166-6851(98)00090-5.

Szecsi, P.B. (1992). The aspartic proteases. Scandinavian journal of clinical and laboratory investigation. Supplementum. 210, 5–22. Published online 1992/01/01.

Toei, M., Saum, R., and Forgac, M. (2010). Regulation and isoform function of the V-ATPases. Biochemistry. 49(23), 4715–4723. DOI: 10.1021/bi100397s.

Tomavo, S., Slomianny, C., Meissner, M., and Carruthers, V.B. (2013). Protein Trafficking through the Endosomal System Prepares Intracellular Parasites for a Home Invasion. PLOS Pathogens. 9(10), e1003629. DOI: 10.1371/journal.ppat.1003629.

Turetzky, J.M., Chu, D.K., Hajagos, B.E., and Bradley, P.J. (2010). Processing and secretion of ROP13: A unique Toxoplasma effector protein. International Journal for Parasitology. 40(9), 1037–1044. DOI: 10.1016/j.ijpara.2010.02.014.

Venugopal, K., Werkmeister, E., Barois, N., Saliou, J.-M., Poncet, A., Huot, L., Sindikubwabo, F., Hakimi, M.A., Langsley, G., Lafont, F., et al. (2017). Dual role of the *Toxoplasma gondii* clathrin adaptor AP1 in the sorting of rhoptry and microneme proteins and in parasite division. PLOS Pathogens. 13(4), e1006331. DOI: 10.1371/journal.ppat.1006331.

Wang, Y., Toei, M., and Forgac, M. (2008). Analysis of the Membrane Topology of Transmembrane Segments in the C-terminal Hydrophobic Domain of the Yeast Vacuolar ATPase Subunit a (Vph1p) by Chemical Modification. Journal of Biological Chemistry. 283(30), 20696–20702. DOI: 10.1074/jbc.M803258200.

Wichroski, M.J., Melton, J.A., Donahue, C.G., Tweten, R.K., and Ward, G.E. (2002). *Clostridium septicum* Alpha-Toxin Is Active against the Parasitic Protozoan *Toxoplasma gondii* and Targets Members of the SAG Family of Glycosylphosphatidylinositol-Anchored Surface Proteins. Infection and Immunity. 70(8), 4353–4361. DOI: 10.1128/iai.70.8.4353-4361.2002.

